# Adult Single-nucleus Neuronal Transcriptomes of Insulin Signaling Mutants Reveal Regulators of Behavior and Learning

**DOI:** 10.1101/2024.02.07.579364

**Authors:** Jonathan St. Ange, Yifei Weng, Morgan E. Stevenson, Rachel Kaletsky, Rebecca S. Moore, Shiyi Zhou, Coleen T. Murphy

**Affiliations:** Lewis Sigler Institute for Integrative Genomics, Princeton University, Princeton, NJ 08544, USA; Department of Molecular Biology, Princeton University, Princeton, NJ 08544, USA

**Author notes:** These authors contributed equally to this work (Jonathan St. Ange, Yifei Weng).

## Abstract

The insulin/insulin-like signaling (IIS) pathway regulates many of *C. elegans’* adult functions, including learning and memory^1^. While whole-worm and tissue-specific transcriptomic analyses have identified IIS targets^2,3^, a higher-resolution single-cell approach is required to identify changes that confer neuron-specific improvements in the long-lived insulin receptor mutant, *daf-2*. To understand how behaviors that are controlled by a small number of neurons change in *daf-2* mutants, we used the deep resolution of single-nucleus RNA sequencing to define each neuron type’s transcriptome in adult wild-type and *daf-2* mutants. First, we found surprising differences between wild-type L4 larval neurons and young adult neurons in chemoreceptor expression, synaptic genes, and learning and memory genes. These Day 1 adult neuron transcriptomes allowed us to identify adult AWC-specific regulators of chemosensory function and to predict neuron-to-neuron peptide/receptor pairs. We then identified gene expression changes that correlate with *daf-2’s* improved cognitive functions, particularly in the AWC sensory neuron that controls learning and associative memory^4^, and used behavioral assays to test their roles in cognitive function. Combining deep single-neuron transcriptomics, genetic manipulation, and behavioral analyses enabled us to identify genes that may function in a single adult neuron to control behavior, including conserved genes that function in learning and memory.

**One-Sentence Summary:** Single-nucleus sequencing of adult wild-type and *daf-2 C. elegans* neurons reveals functionally relevant transcriptional changes, including regulators of chemosensation, learning, and memory.

## Introduction

*C. elegans* is a powerful system for uncovering conserved mechanisms of shared traits, including lifespan regulation, neuronal function, and cognitive behaviors. With 302 adult neurons of only 118 neuron types, individual neurons can have a great impact on behavior. We are particularly interested in the ciliated sensory neuron AWC, which is an amphid odor-sensing neuron that detects the odorants butanone and benzaldehyde^5^ and is required for olfactory learning and memory^1,4^. Many important molecular regulators of individual neuronal functions are highly conserved from worms to mammals. For example, we found that a gain-of-function mutation in a Gαq protein, EGL-30, specifically in the AWC is sufficient to rejuvenate memory in aged worms^6^, and activation of its mammalian homolog, Gnaq, in the hippocampus similarly rescued memory in aged mice^7,8^. Therefore, characterizing the expression of genes in individual neurons may shed light on neuronal function in worms as well as higher organisms. Furthermore, individual chemoreceptors may also determine specific functions of *C. elegans* neurons. For example, the ODR-10 seven-transmembrane chemoreceptor functions in the AWA neurons to detect food and regulate male behaviors^9^. However, unlike the “one olfactory neuron/one receptor” model of the mouse olfactory bulb^7^, *C. elegans* encodes many more than 302 receptors in its genome; whether each receptor is expressed in adult worm neurons or change in specific mutants, and in which neuron they each function, is currently unknown^10–12^. Other neuronal genes such as synaptic and signaling proteins play specific roles in individual neurons, as well.

We are particularly interested in understanding how neuronal behaviors that are controlled by single neurons are regulated by the insulin/IGF signaling (IIS) pathway, which plays an important role in the regulation of longevity, dauer formation, reproduction, and behavior. We previously found that the insulin/IGF receptor mutant, *daf-2*, has extended short-term associative memory^1^, and its long-term memory is extended as well, through increased CREB expression^1,2^. *daf-2* mutants also better maintain motor functions and axon regeneration ability with age^13–15^, and maintain their memory ability with age for twice as long as wild-type worms^16^. We are interested in how the *daf-2* mutation changes the AWC transcriptome, as well as the transcriptomes of other neurons, to change learning ability and allow extended memory function.

Previously, we carried out pan-neuronal transcriptional analysis^2,17^, which is useful for identifying and characterizing previously unknown neuronal genes, but does not have the cellular resolution to determine gene expression in individual neurons. We also performed both whole-worm^3,18^ and neuron-specific RNA sequencing^2,17^ to identify the targets of IIS, and discovered that the neuron-specific targets of *daf-2* are distinct from its whole-body targets^2,3,18^. These neuronal targets regulate phenotypes such as learning, memory, and axon regeneration^2^. However, given the complexity and heterogeneity of the nervous system, bulk sequencing of the entire nervous system cannot detect the location of differential expression, and might omit the differential expression targets that are only present in a subset of neurons. Recent whole-worm single-cell transcriptomic data^19–22^ can distinguish cell types broadly, but lack sufficient resolution and depth in neuronal expression (Extended Data Fig. 1) to answer questions about individual neurons. The CeNGEN database^20^ (Extended Data Fig.1) is an excellent resource for *C. elegans* neuron-specific gene expression for all 118 neuron types from single cell RNA-sequencing of L4 (late larval) stage wild-type hermaphrodites. However, comparing animals of different genetic backgrounds or a different developmental stage or age, even young adults, requires an alternative approach. Previously, Fujiwara et al., found that odor preferences change during development, and adult-specific odor preference is dependent on the FOXO/DAF-16 transcription factor downstream of *daf-2,* and the germline^23^. Therefore, higher-resolution transcriptional profiling of adult neurons from germline-intact animals is necessary to answer these questions.

Single-nucleus RNA-sequencing^24,25^ has become the gold standard for mammalian neuronal sequencing because the neuron’s identity is retained when mRNA is isolated from the nucleus, as opposed to methods where cytoplasmic mRNA from interconnected neurons can be confounded. Here, we adapted this method for *C. elegans* neuronal nuclei to determine the neuronal transcriptomes of Day 1 adult wild-type and *daf-2* mutants. We then used these data to better understand how neuronal transcriptomes shift during the development from L4 to adulthood to allow adult behaviors, what constellation of GPCRs are expressed in adult wild-type worms, and further, how *daf-2* mutation changes individual neuron transcriptomes, resulting in changes in their behaviors. We found that, like behavior, gene expression changes significantly in neurons between L4 and young adult (Day 1), including differential expression of GPCRs in sensory neurons. GPCRs are distributed heterogeneously across the adult *C. elegans* nervous system but are predominantly expressed in chemosensory neurons; in fact, some neurons express hundreds of different GPCRs, suggesting that these neurons must sense and integrate many different sensory inputs. Our data also provide potential neuronal sites of neuropeptide-receptor interactions^12^, which may override the connectome^26^. Analysis of neuron-specific IIS targets revealed significant differences in the regulation of metabolic and longevity genes in specific neurons. Using behavioral experiments, we identified new gene functions in wild-type behavior, as well as neuron-type-specific *daf-2*-regulated genes that play an important role in regulating sensory and cognitive behaviors. In addition to providing an atlas of Day 1 adult wild-type and *daf-2* neuronal gene expression, these data demonstrate that single-nucleus analysis of adult neurons is a powerful tool for understanding neuron-specific changes with the onset of adulthood and in behavioral mutants, and identifying conserved mechanisms of enhanced neuron-specific functions in *daf-2* mutants.

## Results

### Single-nucleus sequencing of adult C. elegans neurons

*C. elegans* develops through a series of four larval stages separated by molts, culminating in reproductively competent adults (Day 1 adults). Similar to Fujibawa et al.^23^, we have observed significant differences in the behavior of L4 larvae and Day 1 adults; in particular, the associative learning ability of L4 larvae is significantly lower than that of adults (Fig. 1a). Because of these differences, data from larval stage neurons, which is the primary existing source of single-cell data available^20^, may not accurately reflect gene expression in adult neurons. Other studies that have carried out adult analyses have used whole animals rather than specifically isolating neurons, meaning that a large proportion of the resulting cells or nuclei are from the germline, which can vastly outnumber other nuclei types. As a result, some whole-worm, germline-intact approaches do not provide sufficient single-neuron-type level information (Extended Data Fig.1). Additionally, approaches that use germlineless mutants might affect the function and transcription of neurons^23^. Finally, to address questions about *daf-2* mutants rather than wild-type animals, we needed to carry out direct experimental comparisons between mutant and wild-type adults.

**Fig. 1.**
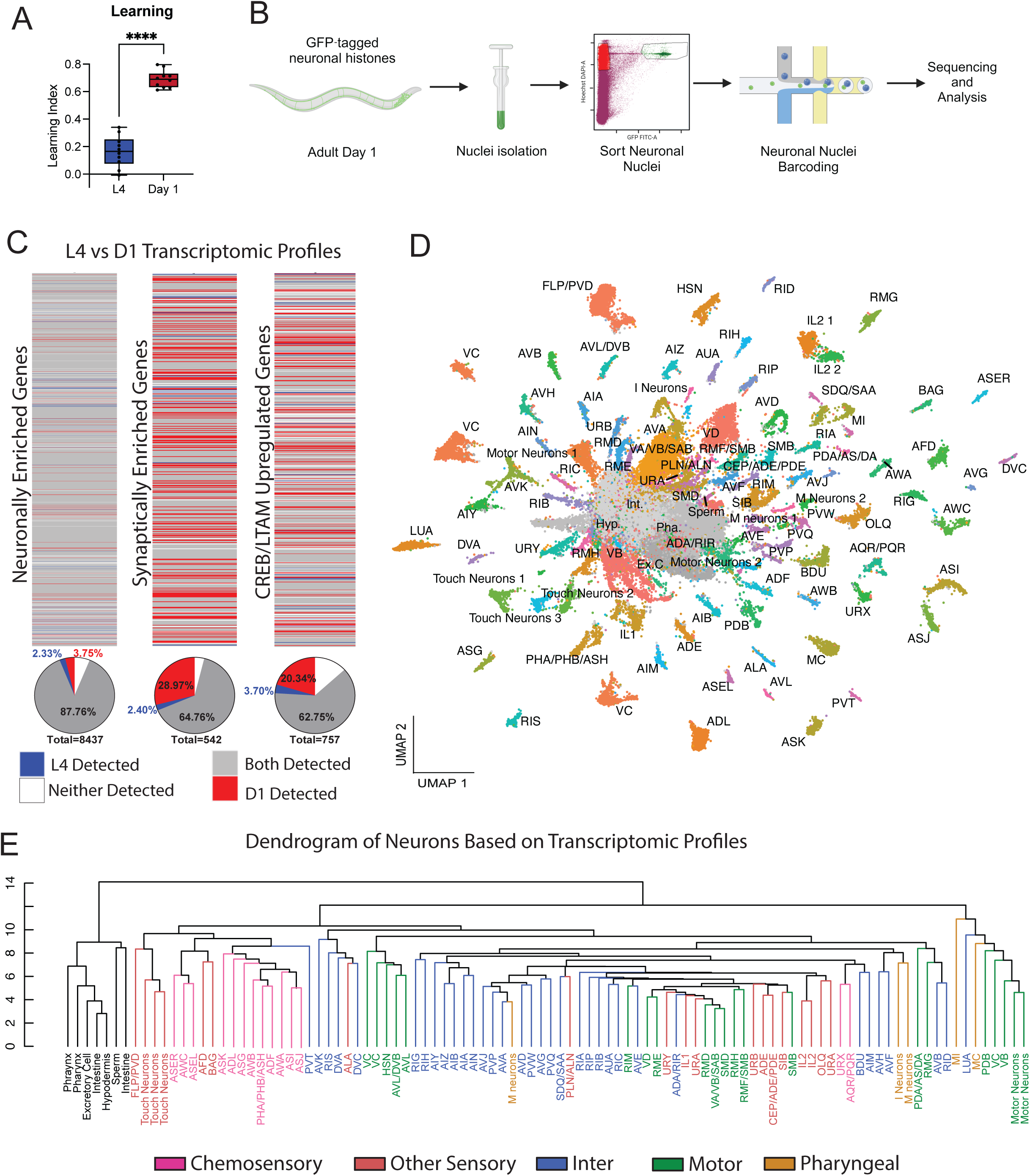
Single Nucleus Sequencing of Adult *C. elegans* Neurons. (**A**) L4 larvae show reduced learning compared to Day 1 adults. Merged data from 2 biological replicates. Each dot represents an individual chemotaxis plate (avg 150 worms/plate). Box plots: center line, median; box range, 25-75^th^ percentiles; whiskers denote minimum-maximum values. Unpaired, two-tailed Student’s t-test. ****p<0.0001. (**B**) An overview of the single-nucleus neuron isolation and sequencing method. *Prgef:his-58::GFP* (Neuronal Nuclei GFP) expressing worms are lysed to release the nuclei, followed by FACS, barcoding using 10X Genomics technology, and Illumina sequencing. Sequencing data were aligned, quality controlled, and clustered using the Louvain Algorithm of unsupervised network clustering of N2 and *daf-2* nuclei. Alignment was performed using Cell Ranger, and all subsequent quality control, normalization, and analyses were done using Seurat. **(C)** Heatmaps comparing L4 transcriptional data (CeNGEN)^20^ to our Day 1 transcriptional data. **(D)** UMAP of all 88,497 nuclei (both wild type and *daf-2*) that passed quality control. **(E)** A hierarchical dendrogram based on gene expression in each Day 1 adult neuron. Neurons are color-coded by functional subtype.

We carried out single-nucleus rather than single-cell RNA sequencing to preserve the connection between cell identity and transcripts. Many *C. elegans* neurons have long processes and intricate synaptic connections, and conventional single-cell RNA sequencing preparation can cause axon breakage, thus losing synapses and allowing RNA to leak out of the cell body. (In fact, we previously took advantage of these neuron disruptions to isolate synaptic fragments and identify synaptically-enriched mRNA transcripts^27^.) By contrast, transcripts inside the nucleus are closely linked with neuron identity. Therefore, we isolated neuronal nuclei from animals expressing a pan-neuronal histone-GFP tag (three biological replicates for each genotype). Both mechanical and chemical lysis were employed to obtain a suspension of nuclei, which were then stained with Hoechst dye. The nuclei were then FACS sorted for both GFP and Hoechst signal (Fig. 1b). Most nuclei maintained their round shape post-FACS and did not leak fluorescence, indicating that the nuclei were not damaged by sorting. RNA from the nuclei was then isolated and sequenced (Methods). After Cell Ranger processing and subsequent ambient RNA removal using SoupX, the samples were assessed for their quality. The number of features (genes), the number of counts (UMIs), and the percentage of mitochondrial transcripts were all assessed, and cutoffs were set to filter out empty droplets and doublets. Each dataset required its own cutoff ranges determined by the mentioned metrics (see Methods); notably, there was no mitochondrial contamination of our samples (Methods). In total, 88,497 neuronal nuclei were obtained after quality control with an average of 756 UMIs/nucleus and 526 genes/nucleus detected (Extended Data Fig. 1). Thus, our sequencing approach using three replicates of wild-type neurons has similar quality metrics to the CeNGEN data that was generated using many samples of overlapping sets of neurons^20^ (Extended Data Fig. 1).

To assess whether our method in fact captured as many or more neuronal transcripts than does conventional single-cell sequencing, we analyzed neuronally-enriched gene transcripts present in our dataset. Our single-nucleus method detected a high percentage of previously-determined neuronal genes (91.5%), suggesting that our method successfully identified neuronal transcripts (Fig. 1c, Tables S1, S2). Thus, despite analyzing slightly fewer neurons (63,565 vs 70,296), our method was able to successfully capture synaptic and neuronal transcripts.

### Neuronal gene expression differs significantly between L4 larvae and Day 1 adults

Because adults differ from L4 larvae in their behavior, we wondered whether their gene expression differs, as well. We examined the differences in expression in synaptically-enriched genes and CREB/long-term associative memory (LTAM) genes in the wild-type subset of our Day 1 adult neuron data and L4 larval neurons from CeNGEN^20^. Our single-nucleus sequencing detected 94% of previously-identified 542 synaptically-localized transcripts^27^ (Table S1), while a comparable single-cell dataset of L4 neurons^20^ only detected 66% of these 542 genes (Fig. 1c). In general, neuronally-enriched genes^17^ are largely present in both datasets (87.8%), suggesting that the highest-expressed genes in neurons are consistent from L4 to Day 1 adults (Fig. 1c, Table S2), although a small number of genes appeared in L4 larvae or Day 1 neurons. However, while 65% of synaptically-enriched genes are shared, many more are detected in Day 1 adult neurons (29.1%) than in L4 larval neurons (2.4%), which could either be due to a difference in single-cell vs single-nucleus approaches, or a shift in expression from L4 larvae to adulthood. By contrast, while 63% of long-term memory genes are shared between L4 and Day 1 neurons, there is a higher representation of these LTAM genes in Day 1 adult neurons (20.4%) that are not present in L4 larval neurons (3.7%). It is unlikely that the higher representation of these genes in Day 1 neurons is a detection artifact, as the L4 larval dataset sampled more than twice the cells (46,627WT nuclei vs 100,955 cells; Extended Data Fig. 1). It is also unlikely that these differences are due to technique differences, because we see other gene categories change, as well; specifically, only 7 of the over 700 LTAM genes are explicitly synaptic, meaning that most of the differences we detect are true differences in expression between stages that are not affected by the difference in cell/nuclei capture approach. Therefore, our data likely reveal shifts in gene expression from the late larval stage to adulthood that may correlate with changes in behavior, further underscoring the need for neuronal transcriptomic data in wild-type adult neurons.

### Identifying neurons from clusters

We next used the Louvain algorithm of unsupervised clustering in the Seurat package^28^ to generate clusters (Fig 1d). To assign cell identities to the clusters, we used a combined systematic and manual curation approach (Methods). This approach included a hypergeometric test along with an area under the curve (AUC) test, followed by manual curation of neuronal anatomies based on the approach by Roux et al.^19^. We used published datasets and reporter expression data, eliminated gene sets assigned to multiple neuronal classifications, and tested different datasets to find consensus on neuronal anatomy (Table S3). These orthogonal approaches increase our confidence in neuron classifications. On average, we found 342 nuclei per annotated neuron cluster (683.5 across two genotypes, wild-type and *daf-2*), on the same order of magnitude as the CeNGEN clusters (549.2 cells/cluster; Extended Data Fig. 1).

109 annotations from our dataset match 118 specific neuron classes (Fig 1c); note that many of the 302 neurons are sorted into specific classes based on left/right pairs and higher-fold symmetry, reducing our expectation of distinct neuron clusters. This classification includes a significant proportion of the neuron population of hermaphrodite *C. elegans*, encompassing 83 of the 87 sensory neurons (95.4%), 68 of the 77 interneurons (88.3%), 101 of the 118 motor neurons (85.6%), and 18 of the 20 pharyngeal neurons (90.0%). Non-neuronal cells, such as intestine, hypodermis, pharynx, sperm, and excretory cells were identifiable and distinguishable from neurons.

### Hierarchical clustering by transcriptome reveals functional relationships

Next, we asked how similar specific neurons are in their gene expression by hierarchical clustering based on gene expression (Fig. 1e). Our dendrogram closely grouped neurons by type, except for a few motor and pharyngeal neurons. This clustering indicates that our sequencing approach captured the cellular identities of different types of neurons and that our neuron identification process was successful. Notably, although neurons arise from specific embryonic lineages, these transcriptomic-based dendrograms are different from anatomy-based lineage dendrograms. For example, URX and AQR/PQR clusters neighbor one another in the transcription-based dendrogram, which is not surprising due to their shared O_2_ and CO_2_ sensing abilities. However, these neurons arise from different lineages: URX develops from the AB cell linage, while AQR and PQR are from the Q cell lineage. Similarly, the AVH and AVF interneurons cluster together due to their similar transcriptomes, which reflects their shared functions as interneurons; however, AVH neurons develop from the AB cell lineage, while AVF neurons are from P1 or W cell lineages. Other neurons cluster together unexpectedly, such as the ALA, DVA, and DVC interneurons, possibly due to their dual functions as interneurons and mechanosensory neurons, or the AIM and BDU interneurons, suggesting that an unidentified functional similarity may exist. The close clustering of the sensory ALN/PLN neurons and SDQ/SAA interneurons might reflect their shared oxygen sensory functions. Thus, transcriptomics of adult neurons provides functional information that may be lacking from existing anatomy and lineage information. These transcriptomic hierarchical clustering results provide an unbiased examination of the relationship of neurons and provide insights into their previously understudied adult functions.

### Neuron-specific transcriptional data provides functional information

To better understand gene expression changes that may correlate with behaviors, we focused on classes of chemoreceptors, which may determine specific functions of *C. elegans* neurons. G protein-coupled receptors (GPCRs) are membrane proteins that serve as receptors for many environmental stimuli. GPCRs bind a wide variety of molecules, from hormones and odors to neurotransmitters^29^ as well as neuropeptides^12^. The number of GPCRs varies greatly between organisms: the *Drosophila* genome encodes around 200 GPCRs^30^, *Ae. aegypti* mosquitos encode 324^31^, mice have 392, and humans express 367 GPCRs^32^, while the *C. elegans* genome encodes over 1300 chemosensory GPCRs alone^33,34^, as well as 153 peptidergic GCPRs, 16 aminergic, 3 muscarinic, and 19 belonging to various other families^34^.

To better understand how chemoreceptors might influence adult behavior, we first determined where chemosensory GPCRs are expressed across all Day 1 adult neurons. Because the exact location of expression of most genes in this large set of GPCRs in adult neurons is unknown, but may contribute to many neuron-specific functions, we first focused on the cell-specific expression of GPCRs. The 1341 *C. elegans* chemosensory (cs) GPCRs have been broken into three super families and six solo families, with 23 families in total^34^; the expression of 375 of the chemosensory GPCRs (28%) have been examined using promoter-GFP reporters^34^, but the location of the remaining 72% and their functions remain unknown, particularly in adult animals. Of the 1341 csGPCRs, we detected 742 in specific neurons at a threshold of 1% detection across a cluster and a normalized average expression value of 0.001, more than doubling the known sites of expression to 56%. We generated a matrix of the percentage of cells expressing each GPCR across all clusters (Table S4); the ADL neuron expresses the most csGPCRs, with 286 in wild-type adult neurons (Fig 2A, S1A, Table S4). Sensory neurons in general express the highest number of csGPCRs, with the ASJ, ASH, ASK, AWA, ASI, ADF, and AWC all expressing dozens to hundreds of receptors (Fig 2a). Chemosensory neurons expressed significantly more csGPCRs than do any other neuron types (Fig 2a); by contrast, peptidergic GPCRs are expressed at a lower and more consistent level across all neuron types (Fig. 2b, Table S5). Our data suggest that many of these sensory neurons and some motor neurons might have the ability to sense information through many receptors simultaneously. In fact, we found no neurons that express only one GPCR, while the vast majority (∼77%) of wild-type neurons express more than 10 GPCRs under normal conditions. GPCR type identification suggested that there was no significant over- or under-representation of any one family, and our dataset identified 50-100% of most families (Extended Data Fig. 3b).

**Fig. 2.**
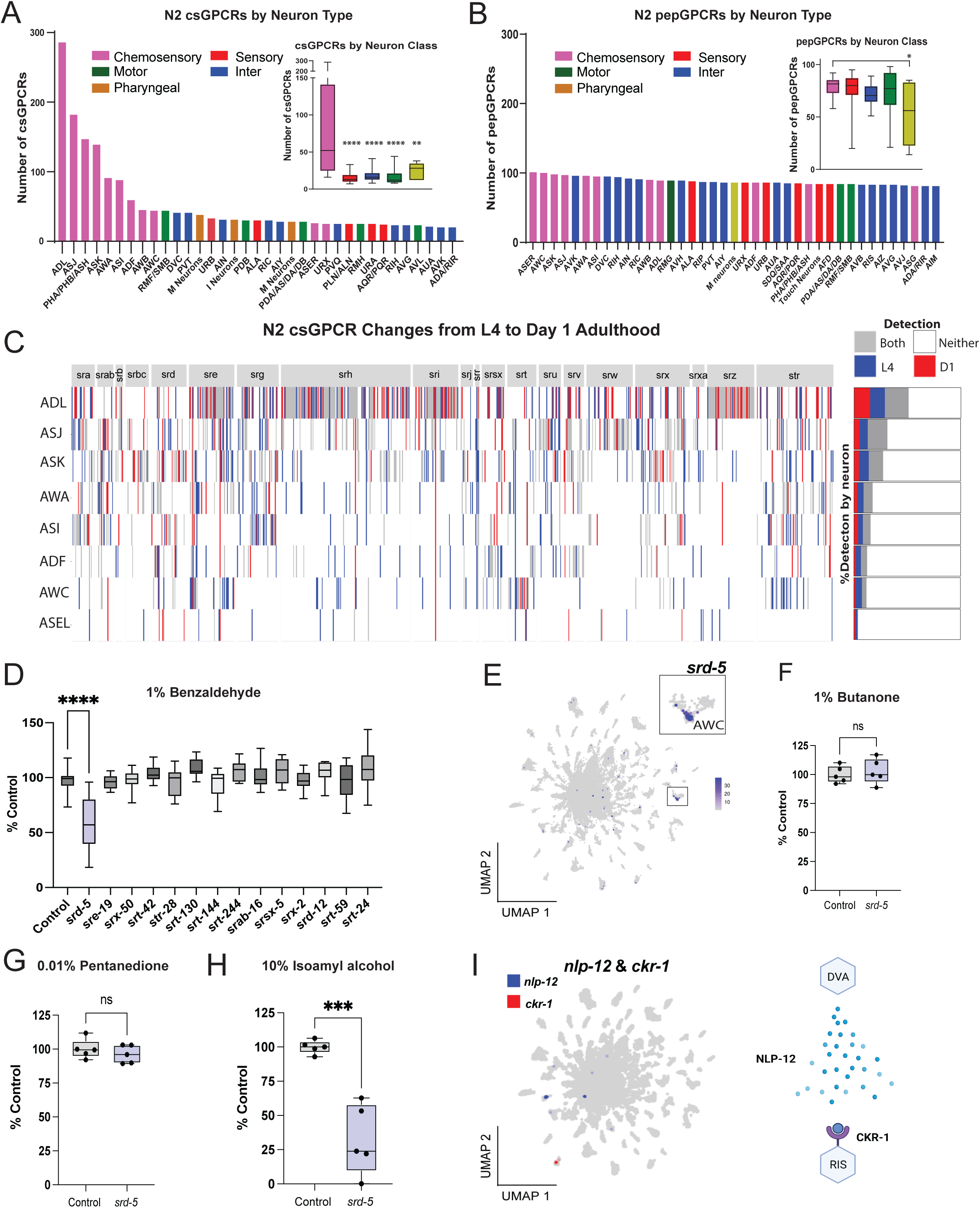
Functional Analysis of GPCRs in the Data Set. (**A**) Chemosensory GPCR (csGPCR) expression in specific neuron classes. Neurons are colored by functional type, as indicated. The top 35 csGPCR-expressing neuron classes are shown. The inset shows csGPCR expression by neuron functional class. (**B**) Peptidergic GPCR (pepGPCR) expression in specific neuron subtypes. Neurons are colored by functional type, as indicated. The top 39 pepGPCR-expressing neuron classes are shown. The inset shows pepGPCR expression by neuron functional class. (**A, B**) Insets: Boxplots: center line, median; box range, 25th–75th percentiles; whiskers denote minimum-maximum values. One-way ANOVA with Tukey’s post-hoc analysis. ****p<0.0001, **p<0.01. *: p<0.05. (**C**) A heatmap of the changes in the csGPCR landscape in several chemosensory neurons from L4 to Day 1. These changes are broken down by GPCR family, and the overall percentage of the changes in each neuron is shown by stacked bar plots. GPCR expression was detected in both L4 and Day 1 neurons (grey), neither dataset (white), only in L4 neurons (CeNGEN)^20^ (blue), or only in Day 1 adult neurons (red). **(D)** Chemotaxis to benzaldehyde upon adult-specific RNAi knockdown of the top Day 1 AWC-only expressed chemoreceptor genes. Pooled data from at least 2 biological replicates. ****p < 0.0001. One-way ANOVA with Bonferroni post-hoc analysis. **(E)** Feature plot of *srd-5* expression across all neuron clusters in our wild type and *daf-2* combined dataset; *srd-5* is primarily expressed in the AWC (inset). (**F-G**) Chemotaxis to butanone (**F)** or pentanedione (**G**) is not affected by *srd-5* knockdown. (**H**) Chemotaxis to isoamyl alcohol is significantly reduced by *srd-5* knockdown. ***: p < 0.001 (F,G,H) Two-tailed t-test, representative figure from 3 biological replicates. (**D, F, G, H**) Neuron-RNAi-sensitized *unc-119p::sid-1* animals were used, and RNAi knockdown was initiated only in adulthood. Boxplots: center line, median; box range, 25th–75th percentiles; whiskers denote minimum-maximum values. **(I)** Feature plot of *nlp-12* and *ckr-1* expression. Schematic based on peptide/receptor interactions described in Beets et al., 2023^12^.

We then asked whether particular families of chemosensory GPCRs are enriched in Day 1 adults compared to L4 larval neurons (Fig. 2c, Table S6). We find that the shifts in gene expression from larvae to adults are distributed across all families. The ciliated amphid ADL neuron, which is involved in several chemosensory roles, including mediating social feeding behaviors and modulating other chemosensory responses, expresses almost 300 csGPCRs, many of which appear to switch “on” in the shift from L4 to Day 1 adults, while others switch “off”. Together, our data suggest that shifts in GPCR expression accompany the transition from L4 larvae to Day 1 of adulthood.

### Functional analysis of GPCRs in the AWC neuron

To test whether chemosensory GPCRs might act in the AWC sensory neuron, which is key for many of the worm’s sensory and olfactory learning abilities^1,4,6^, we knocked down the most highly-expressed candidate csGPCR genes and tested chemotaxis toward odorants that are detected by the AWC neuron^5^. To avoid possible developmental defects, we used RNAi to knock down gene expression in fully-developed neuron-RNAi-sensitive adults for 48 hrs, and subsequently assayed function. First, we tested chemotaxis to benzaldehyde (Fig. 2d), a chemical sensed by both the AWC_on_ and AWC_off_ neurons; although most RNAi treatments had no effect, knockdown of *srd-5,* which we observe is predominately expressed in the AWC neuron (Fig. 2e) reduced chemotaxis to benzaldehyde (Fig. 2d). *srd-5* knockdown did not affect chemotaxis to butanone (Fig. 2f), a chemical sensed only by the AWC_on_, or to pentanedione (AWC_off_) (Fig. 2g). However, chemotaxis to isoamyl alcohol (AWC_on_ and AWC_off_) was reduced by *srd-5* knockdown (Fig. 2h). Our data suggest that the SRD-5 csGPCR is required for detection of benzaldehyde and isoamyl alcohol, odorants sensed by both AWC_on_ and AWC_off,_ but not for sensing of odorants solely by the AWC_on_ (butanone) or AWC_off_ (pentanedione). The identification of SRD-5’s role in AWC-regulated behaviors demonstrates the utility of profiling and functionally testing the chemosensory GPCR landscape in individual neuron types.

### Sequencing data provides locations of potential neuropeptide/receptor interactions

The interactions between neuropeptides (neuropeptide-like proteins, or NLPs, and FMRFamide-related peptides, or FLPs) and their cognate receptors were recently functionally probed in *in vitro* assays to identify potential ligand/receptor pairs, describing a neuropeptidergic network that is independent of the worm’s connectome^10,12^. Randi et al. (2023) used optogenetic measurements to generate a signal propagation atlas, then combined this atlas of extrasynaptic signaling with the potential ligand/receptor pairs from Beets et al., 2023^12^ and gene expression in neurons^20^ to predict pairs of extrasynaptic signaling neuron ligand and receptors. However, these predictions rely on the expression in L4 larval neurons; whether each receptor is expressed in adults, and in which neuron they each function, is currently unknown. Therefore, we used our adult transcriptional data to identify the neuronal sites of potential neuropeptide/receptor interactions; that is, by comparing the neuronal sites of *flp* and *nlp* (Table S7) and neuropeptide receptor transcript expression (Table S8), potential pairwise interactions can be proposed. For example, NLP-58 interacts with TKR-1 and TKR-2^12^; *nlp-58* is expressed in the OLQ and URA neurons, while *tkr-2* is expressed in the ADL neuron, revealing a candidate interaction between the OLQ/URA and ADL neurons (Extended Data Fig. 4c). Although the ADL and OQR are physically connected^35^, the URA-ADL peptidergic interaction may bypass the synaptic connectome. Similarly, we find that NLP-12 is expressed in the DVA interneuron, while its cognate receptor, CKR-1^12^, is expressed in the RIS neuron (Fig. 2j); these neurons are not thought to be directly connected through chemical synapses or gap junctions, but instead may signal via the NLP-12 neuropeptide. Similar maps of adult neuropeptide/receptor interactions can be built by combining these datasets (Table S7, S8), revealing the connections of the adult neuropeptidome that are not obvious from the worm’s connectome.

### Single-nucleus daf-2 sequencing uncovers the heterogeneity of daf-2 regulation in neurons

Our original goal in developing neuronal single-nucleus sequencing (snSeq) approach in adult worms was to characterize expression changes in specific neurons of *daf-2* mutants, particularly those neurons that regulate learning and memory. Therefore, in parallel with wild-type N2, we next performed single-nucleus RNA-sequencing of *daf-2* mutants (three biological replicates) to identify differentially-expressed neuronal IIS targets that might have been masked in our previous pan-neuronal RNA-seq analysis^2,3^. We identified 980 and 695 genes that were up- or down-regulated (log2(Fold-change) > 0.25 or < -0.25, p-adj < 0.05), respectively, in at least one cell type (Fig. 3a, Extended Data Fig. 5a; Table S9). We were surprised to find that, although our single-nucleus dataset is enriched in canonical *daf-2* targets that we had previously identified^2,3^ (Extended Data Fig. 5b), about two-thirds of our dataset consists of previously unidentified *daf-2*-regulated targets (Fig. 3a), indicating that deep, neuron-specific single-nucleus sequencing provides the resolution to identify IIS targets that might have been masked in pan-neuronal, whole-worm, or single-cell analyses.

**Figure 3.**
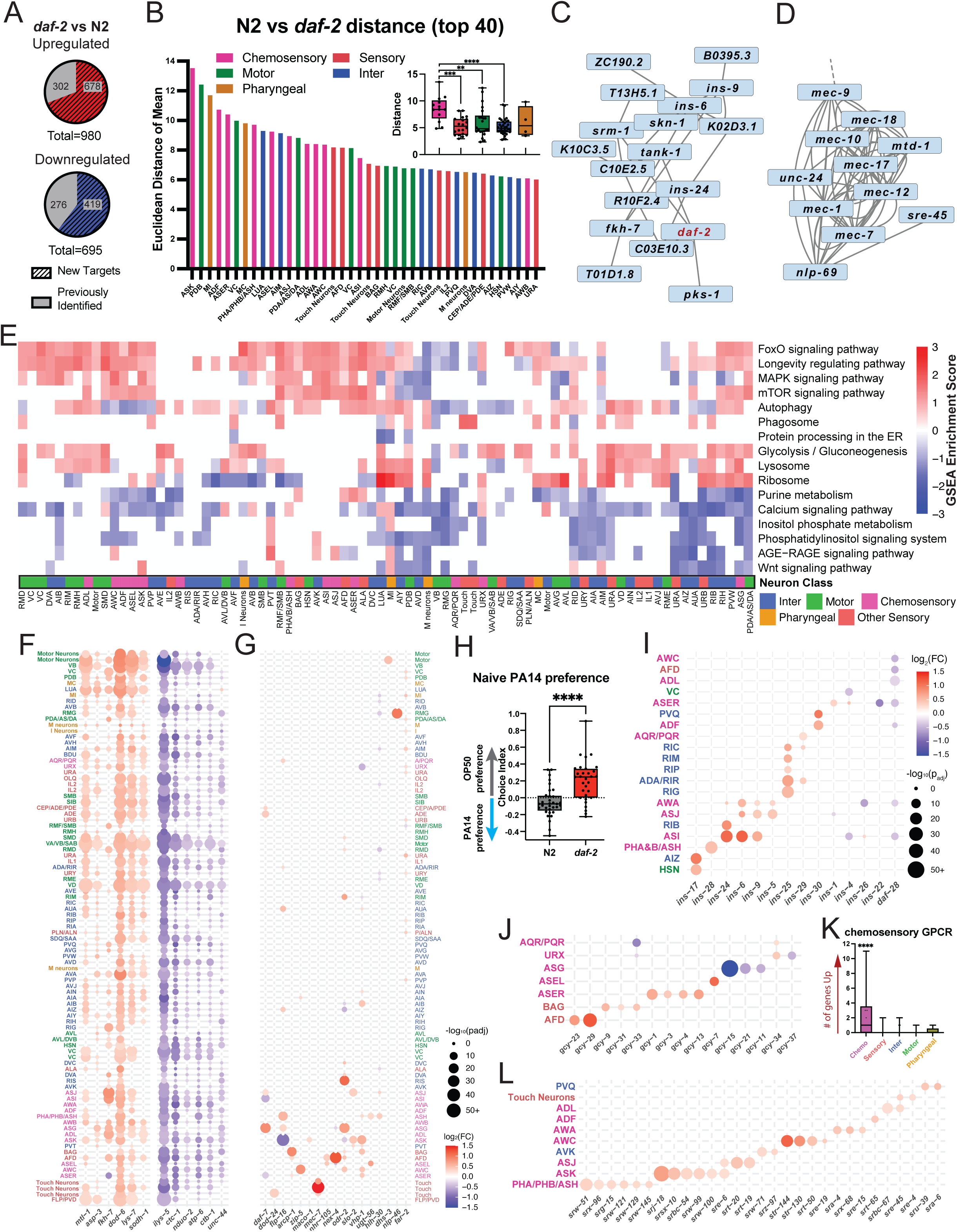
Genes differentially expressed between *daf-2* and N2 vary across cell types. (**A**) Previously and newly identified *daf-2* up-(678) and downregulated (419) genes. (**B**) Euclidean distance of each neuron’s mean *daf-2* vector and mean N2 vector. The top 40 neuron subtypes with the largest distance are shown. The full graph is shown in Extended Data Figure 5C. Inset: average Euclidean distance of each neuron class. Boxplots: center line, median; box range, 25th–75th percentiles; whiskers denote minimum-maximum values. One-way ANOVA with Tukey’s post-hoc analysis. ****p<0.0001, ***p<0.001, **p<0.01. (**C-D**) Gene interaction network based on *daf-2* vs. N2 up-and down-regulated genes. The shorter length of the edges correlates with a stronger connection. (**E**) Heatmap of KEGG pathway GSEA-normalized enrichment scores across different cell types. Empty boxes indicate insufficient overlap between the differentially expressed genes in that cluster and the KEGG pathway gene set to calculate the GSEA enrichment score. Red: pathway is enriched in *daf-2* upregulated genes; (blue) pathway is enriched in *daf-2* downregulated genes. (**F**-**G**) Differential expression of selected genes across neuron subtypes. Colors indicate the direction of differential expression fold-change, and dot size indicates statistical significance from the Wilcoxon Rank Sum Test. (**F**) Differential expressed genes of *daf-2*-regulated targets present in previous studies^2,3,16^. (**G**) Differentially expressed genes newly identified to be differentially expressed in *daf-2* neurons. (**H**) Naïve preference was determined using choice assays to OP50 versus PA14 bacteria. Each dot represents an individual choice assay plate (avg 60 worms/plate). Box plots: center line, median; box range, 25-75^th^ percentiles; whiskers denote minimum-maximum values. Unpaired, two-tailed Student’s t-test. ****p<0.0001. Data from 4 biological replicates are shown. (**I**) Neuron-type expression of insulin-like peptides. (**J**) Neuron-type expression of receptor-type guanylyl cyclases. **K**) Average number of *daf-2*-upregulated GPCRs in each neuronal class. One-way ANOVA with Tukey post-hoc analysis. Box plots: center line, median; box range, 25-75^th^ percentiles; whiskers denote minimum-maximum values. ****p<0.0001. Full comparison shown in the Extended Data Figure 10a. (**L**) Neuron-subtype expression of chemoreceptors that are upregulated in *daf-2* neurons. Full heatmap of GPCR expression shown in Extended Data Figure 9b.

To determine whether some neurons are more affected by the *daf-2* mutation than others, we first calculated each neuron cluster’s Euclidean distance between wild-type and *daf-2* (Table S10; Fig. 3b). In general, we find that sensory neurons, especially chemosensory neurons (ASK, ADF, ADE, ASH, ASJ, etc.) are more distant between *daf-2* and N2 (Fig. 3b inset), consistent with *daf-2’s* notable changes in sensory behaviors^1,36,37^. In particular, gene expression in the ASK neuron, which has roles in multiple types of sensory behaviors, is most different between the *daf-2* mutant and wild-type (Fig. 3b, Fig S5c). This difference is reflected in the PCA plot of ASK (Extended Data Fig. 5d), where the points are separated by genotype more than for neurons with smaller N2 vs. *daf-2* distance, such as the AIB. Some motor neurons also exhibit large differences between N2 and *daf-2,* consistent with extended motor behaviors in aged *daf-2* worms^13,38^. We also examined the correlation between each neuron cluster’s differential expression; similar types of neurons share highly correlated *daf-2*-regulated differential transcriptomic changes, indicating that the *daf-2* mutation effect is neuron-type specific (Extended Data Fig. 6a).

### Global analyses reveal new pathways

We performed gene network analysis by calculating the regulatory strength between all *daf-2* significantly differentially-expressed genes using tree-based ensemble methods (GENIE3^39^) to investigate the co-regulation and interaction of these differentially expressed genes in the nervous system (Extended Data Fig. 7). Several insulin-signaling-related genes form an interaction network with *daf-2*, including insulin-like peptide genes (*ins-6*, *ins-9*, *ins-24*), a forkhead transcription factor (*fkh-7)*, and the *skn-1/Nrf* transcription factor (Fig. 3c). Interestingly, there are also other genes in this hub that have not been previously associated with insulin signaling, such as the NAD^+^ ADP-ribosyltransferase *tank-1/pme-5*, the fatty acid (polyketide) synthase *pks-1*, the serpentine receptor *srm-1*, a protein tyrosine phosphatase receptor (*T13H5.1*), an elongation factor (*K10C3.5*), and uncharacterized genes. PME-5 is involved in the *C. elegans* apoptosis DNA damage response pathway^40^. *K10C3.5* expression is induced by UV exposure in *daf-2* mutants and its induction is DAF-16-dependent^41^. *K02D3.1* is significantly downregulated in *daf-16* mutants^42^ and upregulated in *skn-1*^43^ mutant background. Our network analysis also suggested that mechanosensory genes (e.g., *mec-1*, *mec-7, mec-9, mec-10*, *mec-12*, *mec-17, mec-18*, *mtd-1, unc-24, nlp-69,* as well as an epsilon class serpentine receptor (*sre-45*)) are differentially expressed in *daf-2* mutants (Fig. 3d). Together, these data suggest that there are *daf-2*-regulated neuronal functions that have not yet been identified, and serve as a resource for building hypotheses towards identifying and characterizing these functions.

KEGG analysis of *daf-2*-regulated genes revealed that FOXO, MAPK, mTOR signaling, autophagy, and other longevity-related pathways are upregulated in most neurons, consistent with canonical IIS/FOXO pathway functions (Fig. 3e, S8a; Table S11). By contrast, purine metabolism, calcium signaling, and inositol-phosphate metabolism pathways – all associated with neuronal activity - were downregulated in most neurons. Although decreased activity seems surprising, neuronal hyperactivity has been linked with cognitive decline and neurodegeneration^44,45^; downregulation of calcium and inositol signaling genes might reduce hyperactivation with age. The AGE-RAGE signaling pathway, which is involved in binding Advanced Glycation End damage products that are produced by the Maillard reaction and is associated with diabetes^46^, is also downregulated in select neurons. Some pathways are more heterogenous, such as ribosomal genes (*rps* and *rpl*), which were upregulated in *daf-2* in some neurons (e.g., LUA, AIY, AVL) and downregulated in others (e.g., RIC, AVB, RMF/SMB). Interestingly, ribosomal biogenesis is universally downregulated in longevity mutants to reduce energy expenditure^47,48^, but our data suggest this global downregulation could have neuron-specific exceptions.

### snSeq analysis reveals genes changed in single neurons

Canonical *daf-2*-upregulated (Class 1) genes, such as *mtl-1, fkh-7, dod-6,* and *lys-7*, and downregulated (Class 2) genes (*ilys-5, ctc-1, nduo-2*) were differentially expressed across many neurons (Fig. 3f), consistent with our previous pan-neuronal results^2^. Although the forkhead transcription factor *fkh-7* was predominantly upregulated in *daf-2* neurons, it was downregulated in a single neuron, the ASER. By contrast, newly-identified *daf-2*-regulated genes, such as *flp-16, zip-5, maco-1, mec-7, nex-4, odr-2, slo-2, nlp-46,* and *far-2*, were differentially expressed only in a few neurons (Fig. 3g). In fact, when we examined how genes are differentially expressed between *daf-2* and wild-type across neuron types, we found that over half were only differentially expressed in a single neuron, and over 98% of genes were differentially expressed in fewer than 30 neuron types (Extended Data Fig. 8b). That is, the expression of a particular gene may be different in only a handful of neurons, making it difficult to discover from whole-worm transcriptomics, tissue-specific bulk RNA-seq, or shallow single-cell RNA sequencing data – even if that gene expression difference in a particular neuron is critical for a functional difference. Others are significantly upregulated in one neuron but downregulated in another, thus its net differential expression in previous bulk analyses might be negligible. For example, *daf-7* is upregulated by *daf-2* in the ASJ, ASG, and ADE neurons, but downregulated in the ASI and ASER neurons. Because increased *daf-7* expression in either ASJ^49^ or ASI^50^ is sufficient to induce a switch from attraction to PA14 to avoidance, we wondered whether *daf-2* mutants might have altered chemotaxis to PA14. Indeed, *daf-2* mutants show a higher naïve avoidance of PA14 than do wild-type worms (Fig. 3h).

These newly-identified, *daf-2*-regulated genes regulate many aspects of the nervous system. For example, ODR-2 regulates odor sensation and pheromone imprinting^5,51^, *mec-7* encodes a beta-tubulin gene that regulates mechanosensory neuron development and touch sensitivity of worms^52^, and ZIP-5 regulates axon regeneration^15^ and pathogen-induced pheromone responses^53^. The extended health and behaviors of *daf*-*2* animals may result from such neuron-specific gene changes that were previously masked.

Next, we examined how specific gene families are regulated in individual neurons by *daf-2*. First, we found that insulin-like peptides were differentially expressed in *daf-2* mutants in specific neurons, most of which are chemosensory or head interneurons (Fig. 3i); *ins-17* (AIZ), *ins-24* (ASI) and *ins-6* (ASI) show particularly strong differential expression. Similarly, receptor-type guanylyl cyclases were more differentially expressed specifically in the ASEL (*gcy-7*), ASER (*gcy-1, -3, -4,* and *gcy-13*), ASG (*gcy-15*), and AFD (*gcy-23* and *gcy-29*) sensory neurons in *daf-2* mutants (Fig 3j).

To identify *daf-2*-differentially expressed GPCRs, we plotted the expression (Extended Data Fig. 9a) and differential expression (Extended Data Fig. 9b) of chemosensory GPCRs, peptidergic GPCRs, and other GPCRs in *daf-2* neurons. While GCPRs are widely expressed (Extended Data Fig. 9a), they are selectively differentially expressed only in a subset of neurons (Extended Data Fig. 9b). In general, more chemosensory GPCRs are up-than downregulated in *daf-2* mutants (Extended Data Fig. 10a-b), which is also true for peptidergic GPCRs (Extended Data Fig. 10a-b). Chemosensory neurons on average have the greatest number of upregulated GPCRs in *daf-2* mutants (about 5 per neuron), significantly higher than other neuron types (< one) (Fig. 3k). Chemosensory neurons are the site of 76% of all upregulated chemosensory GPCRs (Extended Data Fig. 10b); many chemosensory GPCRs were upregulated in *daf-2* in a single sensory neuron type, e.g., *srj-18* in the ASK, and *str-144*, *str-130,* and *str-50* in the AWC neuron (Fig. 3i). The fact that most of these GPCRs are upregulated in *daf-2* mutants (Fig. 3l) suggests that *daf-2* neurons may have a unique advantage over wild type in chemosensory-related behaviors. Other GPCRs are also differentially expressed in *daf-2* mutants, but in various neuron types (Extended Data Fig. 9b). Some of these GPCRs have important neurotransmitter signaling functions, including dopamine receptor *dop-3* (ASK, AVK) and glutamate receptor *mgl-3* (ASK), serotonin receptor *ser-4* (ASI and AWA), octopamine receptors *ser-3* (AVK) and *ser-6* (ADE), and tyramine receptors *ser-2* (DVA) and *tyra-2* (AWC) (Fig. 9a).

### Genes that normally decline with age are upregulated in daf-2 neurons

In addition to their greatly extended lifespan, *daf-2* mutants have improved and extended cognitive and neuronal behaviors^1,6,15,36^, some of which may depend on gene expression changes in individual neurons. We previously used pan-neuronal transcriptional profiling to identify genes whose expression is high in young wild-type neurons and declines with age^16^. These genes are enriched in neuronal and synaptic function^16,54^, and their downregulation with age correlates with behavioral deficits that rise with age. With the exception of the ubiquitously-downregulated lysozyme *ilys-5* (Extended Data Fig. 10c) and the dietary restriction overexpressed *droe-4* (AWA), most of the genes in this category are upregulated in *daf-2* mutants (Fig. 4a). While the ubiquitously upregulated genes include collagens^17^, the DAF-16 target *dct-16*^55^, a nematode allergen (*npa-1*), and the *cpr-*9 and *asp-3* proteases, the majority of these age-downregulated genes are upregulated only in a small subset of *daf-2* neurons. For example, several *mec-* genes are specifically upregulated in the mechanosensory touch neurons, insulins, neuropeptides (*nlp-*, *flp-*), neuropeptide receptors (*pdf-1, pdf-2, tkr-3*), chemosensory genes (*odr-2, hot-1*), ion channels, transmembrane transporters, and synaptic transmission genes are upregulated in specific neurons (Fig. 4a).

**Figure 4.**
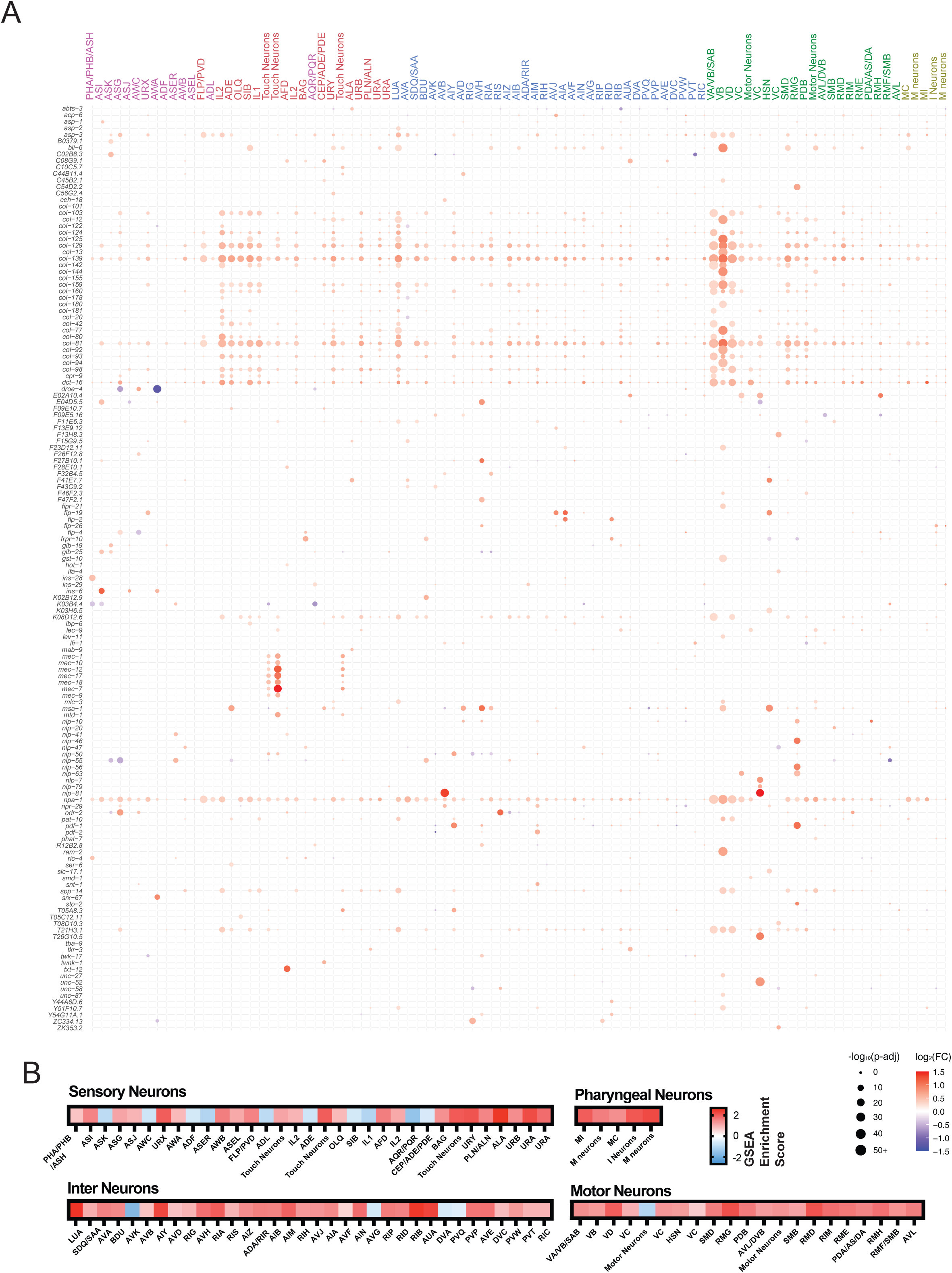
*daf-2* exhibits changes in aging gene expression in specific neurons. (**A**) Expression map of known neuronal genes that are downregulated in wild-type neurons with age^16^ that are significantly upregulated in at least one neuron type in Day 1 *daf-2* neurons. Neurons are color-coded based on their functional class. (**B**) GSEA enrichment score of genes that are downregulated with age in wild-type neurons that differentially change in *daf-2* neuron subtypes. Each neuron’s *daf-2* differentially expressed genes were generated into a ranked list from the most upregulated to downregulated in *daf-2*. GSEA analysis was performed by comparing *daf-2* neuron ranked lists to genes that are downregulated with age in wild-type neurons. Red: enrichment of genes upregulated in *daf-2*; blue: enrichment of genes downregulated in *daf-2*.

The genes that decline with age in wild-type neurons^16^ are enriched in all of *daf-2’s* neurons (Fig. 4b), although some sensory neurons show lower GSEA (gene set enrichment) scores; this may be due to the relatively low representation of sensory neurons in the bulk sequencing results due to dissociation and sorting difficulties, or because of their unique transcriptomes, while inter, pharyngeal, and motor neurons are more homogenous. Unbiased oligo enrichment analysis of the 1kb upstream region of the *daf-2* upregulated genes revealed the FOXO/DAF-16 binding element (DBE) (GTAAACA), suggesting that many of the *daf-2*-regulated genes are DAF-16 targets^56^ (Extended Data Fig. 10D, Table S12).

### AWC-specific daf-2 targets function in butanone sensation and associative memory

The AWCs are two chemosensory neurons that are collectively responsible for butanone, benzaldehyde, and isoamyl alcohol sensation, and are required for butanone appetitive associative learning and memory enhancement in the *daf-2* mutants^1,4–6^. The AWC neuronal transcriptome is significantly enriched in the expression of ion transport, protein kinase, and ATP and GTP binding activities compared with all neurons, indicating its role in signal transduction (Fig. 5a).

**Figure 5.**
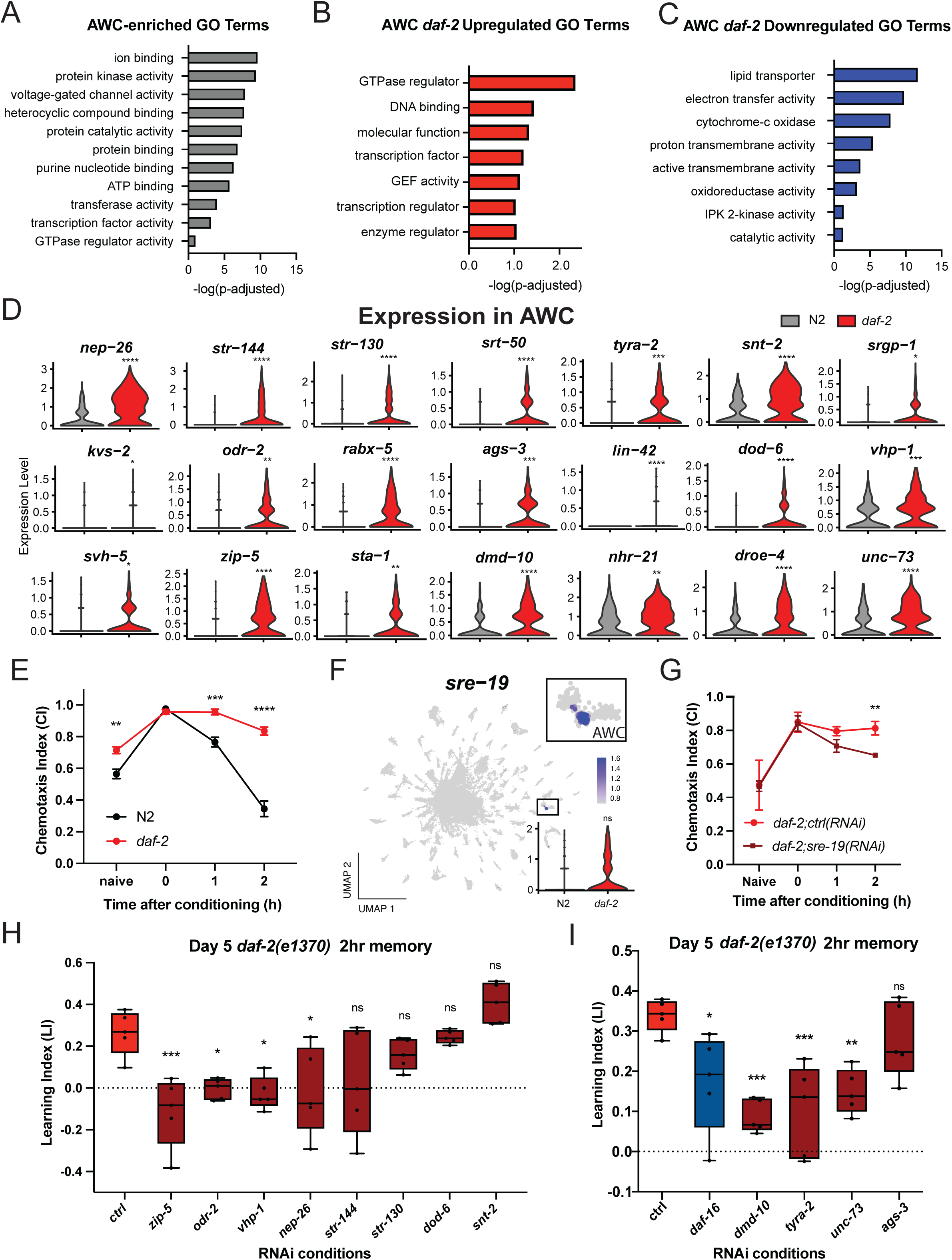
AWC-specific *daf-2*-regulated genes function in chemosensation and cognitive functions. (**A**) Significant Gene Ontology (GO) terms for AWC-enriched genes, which are genes significantly upregulated in the combined dataset of wild-type and *daf-2* AWC neurons compared with the total transcriptome (log_2_[Fold-change (AWC/total)] > 0.25, p-adj < 0.05, Wilcoxon Rank Sum Test). (**B**) GO terms of AWC-specific *daf-2*-upregulated genes (*daf-2* vs N2 log_2_(fold-change) > 0.25, p-adj <0.05, Wilcoxon Rank Sum Test). (**C**) GO terms of AWC-specific *daf-2*-downregulated genes (*daf-2* vs N2 log_2_(fold-change) < -0.25, p-adj <0.05, Wilcoxon Rank Sum Test). GO Terms generated using WormCat 2.0^67^. P-values of GO Terms were Bonferroni corrected. (**D**) Expression distribution of AWC-upregulated *daf-2* genes. Expression level density of (grey) N2 or (red) *daf-2* AWC cells. Adjusted p-values from Wilcoxon Rank Sum test. *p<0.05. **p<0.01. ***p<0.001. ****p<0.0001. Exact p-values are shown in Supplemental Table 7. (**E**) *daf-2* mutants have significantly better short-term associative memory (STAM) with age at both 1hr and 2hr post-training compared to wild-type worms (Day 6 shown here). Representative figure from 3 biological replicates. Error bar: SEM. ****p < 0.0001, Two-way ANOVA with Sidak’s post-hoc analysis. Error bars: SEM. (**F**) Feature Plot expression level of *sre-19*. The scale bar is shown for the SCT normalized expression level. Inset 1: *sre-19* expression in the AWC. Inset 2: *sre-19* expression distribution in N2 and *daf-2* AWC neurons. (**G**) STAM of Day 5 neuronal-RNAi sensitive *daf-2* worms treated with adult-specific *sre-19* RNAi as adults. Representative figure from 3 biological replicates. **p = 0.0098. Two-way ANOVA with Sidak’s post-hoc analysis. Error bars: SEM. (**H**-**I**) 2-hour STAM of Day 5 neuronal RNAi sensitive *daf-2* worms treated with either adult-specific Control RNAi (empty L4440 vector) or candidate gene RNAi. *daf-16* RNAi was used as a positive control. Each dot represents an individual chemotaxis plate (avg 150 worms/plate). Representative image of 2 biological replicates. One-way ANOVA with Dunnett’s post-hoc analysis. Boxplots: center line, median; box range, 25th–75th percentiles; whiskers denote minimum-maximum values. *: p <0.05, **p<0.01, ***p <0.001, ****p<0.0001.

In *daf-2* mutants, the AWC exhibits significant differences from wild type in expression of GTPases, GEFs, and transcription factors, and transmembrane transport and oxidoreductase activities (Fig. 5b-c; Table S13). Upregulated genes include neprilysin (*nep-26*), several Seven-transmembrane, serpentine, and tyramine receptors (*str-144, str-130, srt-50, tyra-2*), genes encoding synaptic proteins (synaptotagmin 2/*snt-2*), axon guidance proteins (Slit-Robo GAP homolog 1/*srgp-1*), a potassium voltage-sensitive channel (*kvs-2*), and the ODR-2 GPI-linked signaling protein, as well as other signaling proteins (activator of G protein signaling/*ags-3*, RAB exchange factor/*rabx-5,* Rho GEF Trio*/unc-73*), the *lin-42* miRNA regulator, DAF-16 target *dod-6,* VH1-related genes (*vhp-1, svh-5*), transcription factors (bZIP/*zip-5*, STAT/*sta-1*, Doublesex/*dmd-10,* and NHR-21), and the Dietary Restriction OverExpressed gene *droe-4* (Fig. 5d; Table S9).

To understand whether genes differentially expressed in AWC neurons might affect behavior, we carried out functional analyses. *daf-2* worms have significantly higher butanone short-term associative memory than wild type in both young and older animals (Fig. 5e, Extended Data Fig. 11b), and the AWC is the butanone olfactory learning and memory center^6,57^. The expression of one serpentine receptor, *sre-19*, is confined to the AWC neurons (Fig. 5f); therefore, we tested whether its knockdown would affect *daf-2’* improved memory with age. (To avoid any effects during the development that might confound interpretation of adult roles of the genes, we use RNAi to knock down gene function solely in adulthood, and we performed the STAM assay on Day 5 because, at this time, *daf-2* worms have intact memory, but wild-type worms have reduced memory, thus if the *daf-2*-upregulated genes are required for *daf-2*’s memory improvement, knocking them down should reduce memory.) We knocked down *sre-19* in *daf-2* adults and found that this receptor, although it did not affect motility, or the ability to sense butanone (naïve chemotaxis, Extended Data Fig. 11c-d), was indeed required for aged *daf-2*’s extended short-term memory (Fig. 5g). We also selected top *daf-2*-upregulated AWC-expressed genes (Fig. 5d) whose RNA interference strains^58^ were available to test for their effects on *daf-2’s* extended memory (Fig. 5h, i). Although not all of the top *daf-2*-upregulated AWC genes had an effect, seven of the 12 tested candidates (*nep-26, vhp-1, odr-2, zip-5, dmd-10, unc-73,* and *tyra-2*) are required for extended short-term memory in aged *daf-2* worms. We observed no differences in motility, chemotaxis, or other behaviors, suggesting that the knockdown of these genes did not affect non-AWC cells. These genes, as well as others in the *daf-2* AWC-upregulated set, likely play important roles in cellular signaling and regulation during the memory formation and storage process.

## Discussion

Here we have used single-nucleus sequencing (snSeq) to transcriptionally characterize each of *C. elegans*’ adult neurons. Because transcripts in isolated nuclei are not subject to loss due to close connections with other neurons or disruption during preparation, single-nucleus RNA sequencing is particularly useful for neuronal transcriptional analyses. Our results indicate that single-nucleus sequencing can identify targets in specific neurons that were masked in whole-worm or tissue-specific sequencing efforts, revealing new behavioral regulators. Furthermore, because L4 larvae differ in neuron-regulated behaviors compared to Day 1 adults, the identification of transcriptional differences in neurons can help us uncover the causes of these differences. For example, we find that L4 larvae cannot carry out standard butanone associative learning behavior; Day 1 animals may need to perform similar behaviors, while the transient state of L4, which just needs to make it to adulthood, may be reflected in its transcriptome. In fact, most of the genes that we had previously identified in studies of learning and memory are only expressed after the transition from L4 to adult neurons. Moreover, the transcriptomes of adult neurons reveal potential functional similarities that are not obvious from lineage or morphological data.

Transcriptional changes of a single gene within an individual neuron can be detected using snSeq, allowing us to better describe the expression profile of *C. elegans’* large family of GPCRs. While the expression of 29% of these GPCRs was already known, our results have determined the site of expression of an additional 554 GPCRs. Remarkably, most neurons express not just one, but multiple GPCRs, and some sensory neurons express dozens to hundreds of these receptors, suggesting that these neurons have at least the potential to sense many inputs simultaneously. For example, the ciliated amphid sensory neuron ADL expresses more than three hundred GPCRs; how these receptors respond to and integrate such a vast number of inputs, and change in different mutant backgrounds, will be interesting to decipher. We also used this information to identify and test candidate GPCRs for possible AWC-neuron-related chemosensory functions, and to identify potential neuronal sites of candidate neuropeptide-receptor interactions. The SRD-5 chemosensory GPCR, for example, is necessary specifically for benzaldehyde and isoamyl alcohol chemotaxis, but not for other AWC sensory functions. Our data also provide expression data for candidate neuropeptide-receptor interactions, helping us to better understand the adult peptidergic signaling connectome.

Single-nucleus sequencing is also a powerful tool to identify differential gene expression changes in single neurons of mutants, which cannot be inferred from existing datasets of wild-type neurons. Changes in single neurons that would have been masked in whole worm, pan-neuronal, and even single-cell analyses are identifiable in our snSeq data. For example, we found that over half of *daf-2*-regulated genes were only differentially expressed in one neuron type and were identified as neuronal *daf-2* targets for the first time here. Other genes have surprisingly restricted expression changes; for example, the serpentine receptor *sre-19* was only expressed in the AWC neuron and is required for *daf-2’s* ability to extend short-term memory. Likewise, insulin-like peptides have restricted expression; *ins-4* was downregulated only in ASI and VC neurons in *daf-2* mutants, which corresponds to its function in pheromone-mediated learned avoidance change^59^, and *ins-6* was upregulated in *daf-2* ASI neurons, which correlates with its function in aversive memory formation in ASI^60^. We also found that guanylyl cyclases, which function in salt chemotaxis, olfaction, CO_2_ sensation, and thermotaxis, were differentially expressed in *daf-2* only in the sensory neurons mediating these responses, correlating with the altered behaviors seen in *daf-2* mutants^61,62^. Most of these gene expression changes were not obvious in bulk RNA sequencing, likely due to fact that the changes are restricted to only a single or small number of neurons, or in some cases, might have been masked by decreases in other cells. Thus, single-nucleus sequencing is a powerful tool for identifying genes that only change in a subset of neurons and would be masked in tissue-specific or whole-worm analyses.

We also detected new AWC-specific *daf-2*-upregulated genes that were required for AWC-mediated learning and memory. Of these genes, several were previously known to have neuronal functions, but had not been previously associated with AWC or *daf-2* phenotypes. For example, ZIP-5 was previously shown to function in axon regeneration of *daf-2* worms^15^, but here we found that *zip-5* is upregulated in *daf-2* and is required for short-term associative olfactory memory. ODR-2 is important for AWC chemosensory functions^63^ and has been implicated in odor imprinting memory^51^, but was not previously known to be required for *daf-2’s* extended memory (Fig. 3f, 5h). The transcription factor *dmd-10* was previously shown to be required for ASH-mediated osmolarity avoidance^64^, but its requirement in the AWC neuron had not been previously shown. Likewise, the TYRA-2 tyramine receptor that is required for imprinted memory of pathogen avoidance^65^, is also required for the extended memory ability of *daf-2* worms (Fig 5d, i). The Rho GEF UNC-73 has been shown to regulate axon guidance, motility, and learning in *C. elegans,* and its homolog Trio is required for normal spatial learning in mice, but its connection with the insulin signaling pathway was not previously described^6,66^. The role of UNC-73 in *daf-2’s* memory extension echoes our recent finding that axon guidance factors are required for the extended memory function of activated G_α_q proteins in aged animals^8^. The upregulation of these genes may promote structural and functional integrity of the neuron and facilitate metabolic processes and signal transport, improving these behaviors relative to wild type, and maintaining these behaviors with age.

Our data further describe the expression landscape of these IIS differentially-expressed genes and link them to new roles downstream of *daf-2* to regulate learning and memory. Together, our results provide new deep transcriptional data for individual adult *C. elegans* wild-type neurons, and suggest that we can use single-nucleus sequencing to identify cell-type specific changes in adult mutants to uncover new conserved candidates that regulate sensory and cognitive behaviors.

---

**Extended Data Figure 1.**
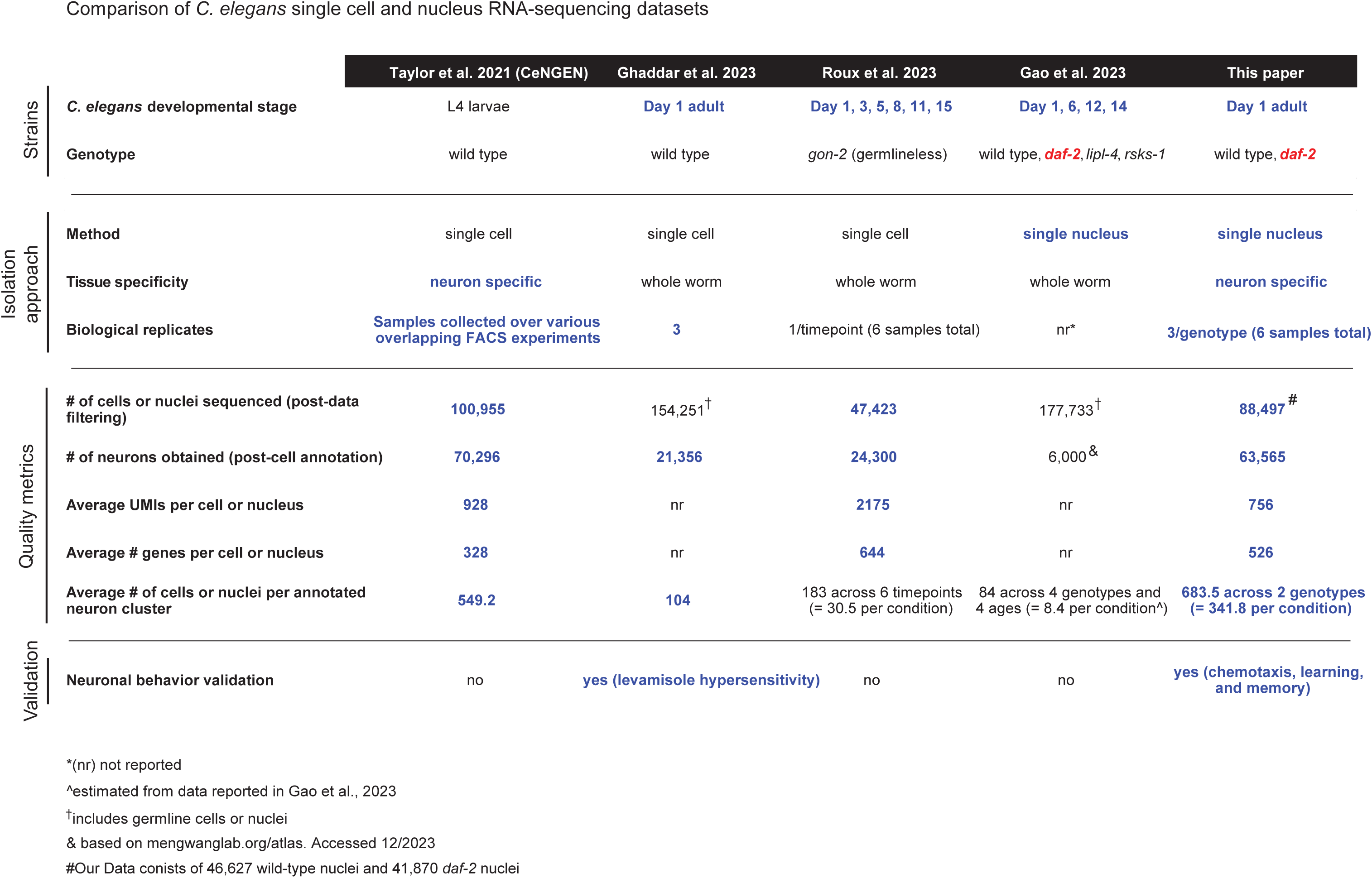
Table detailing methods and metrics of different RNA-sequencing datasets. The isolation approach of single nucleus sequencing provided many benefits, and biological replicates strengthen our confidence in the data. To perform a neuronal analysis, thousands of neurons are required; therefore, a large number of cells/nuclei for any specific neuron type are very important. Finally, functional validation of RNA-sequencing data demonstrates the power of a given method.

**Extended Data Figure 2.**
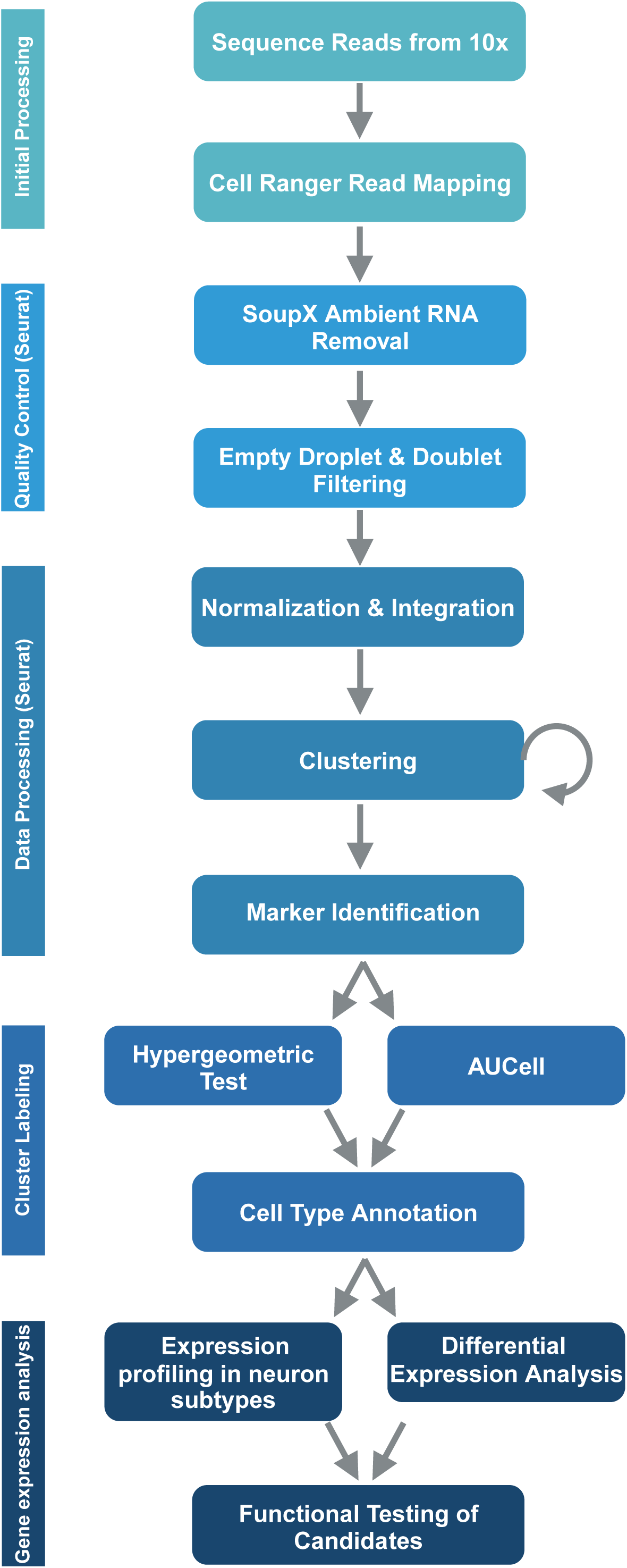
RNA-sequencing data analysis pipeline. 3 biological replicates of both wild-type animals and *daf-2* mutant animals were collected, and following nuclei isolation and library generation, reads were sequenced using 10x. Using Cell Ranger version 7.0.1 with default settings, the reads were mapped to the Ensembl *C. elegans* N2 genome. Next, Cell Ranger output files were fed directly into the SoupX function for ambient RNA removal. Individual samples were made into Seurat objects, and then the distribution of features and counts allowed us to filter out empty droplets and doublets. Data was merged into a single Seurat object and then normalized using SCTransform and integrated. Clustering was an iterative process where multiple amounts of principal components (from 100-200) were tested, as well as multiple clustering resolutions (from 0.5-2) before we settled on 150PCs at a resolution of 1. Markers were then identified with the ‘FindAllMarkers’ function in Seurat, and these markers were used for a hypergeometric test and an AUCell test, as detailed in Roux et al., 2023^19^. Next, we manually compared the two tests looking at their agreement to give clusters their final labels with high confidence. Finally, we were able to profile the expression of genes across individual neurons, perform differential expression analysis across our two genotypes, and functionally test candidates that arose from our data.

**Extended Data Figure 3.**
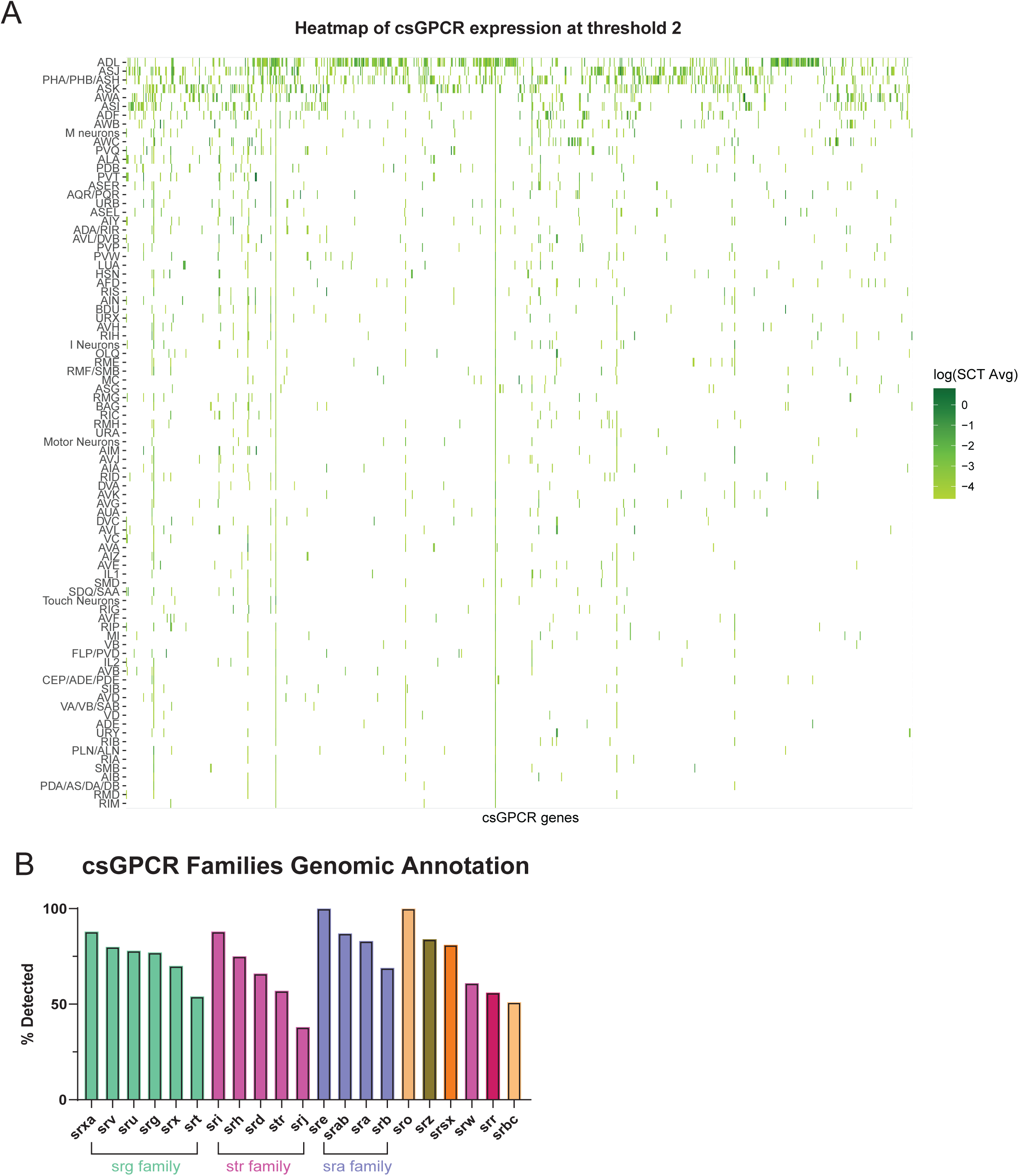
Extra information on GPCR expression across our dataset. (**A**) A heatmap of chemosensory GPCRs (csGPCRs) across all neuron classes at a detection threshold of 2. (**B**) Percentage of GPCRs detected in the dataset organized by family. The percentage is based on the number of members of the family detected divided by the genomic annotation_65_.

**Extended Data Figure 4.**
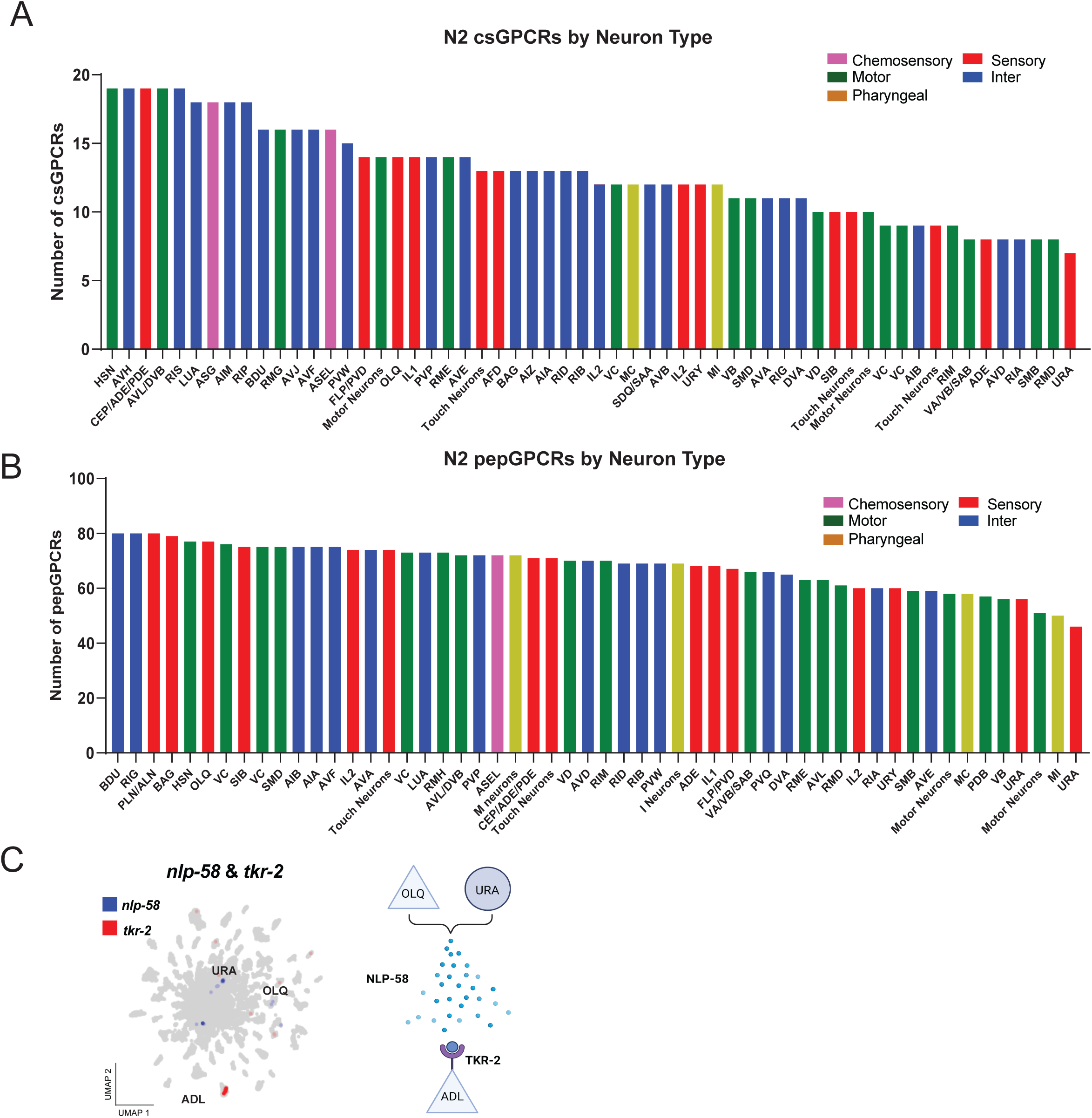
Neuron type analysis of GPCRs. (**A**) Bar graph showing the number of csGPCRs by neuron type. These neuron subclasses are the remainder not shown in Figure 2A. (**B**) Bar graph showing the number of peptidergic GPCRs (pepGPCRs) by neuron type. These neuron subclasses are the remainder not shown in Figure 2B. (**C**) A schematic of a predicted neuronal signaling interaction based on a neuropeptide and neuropeptide receptor interaction characterized in Beets et al_12_.

**Extended Data Figure 5.**
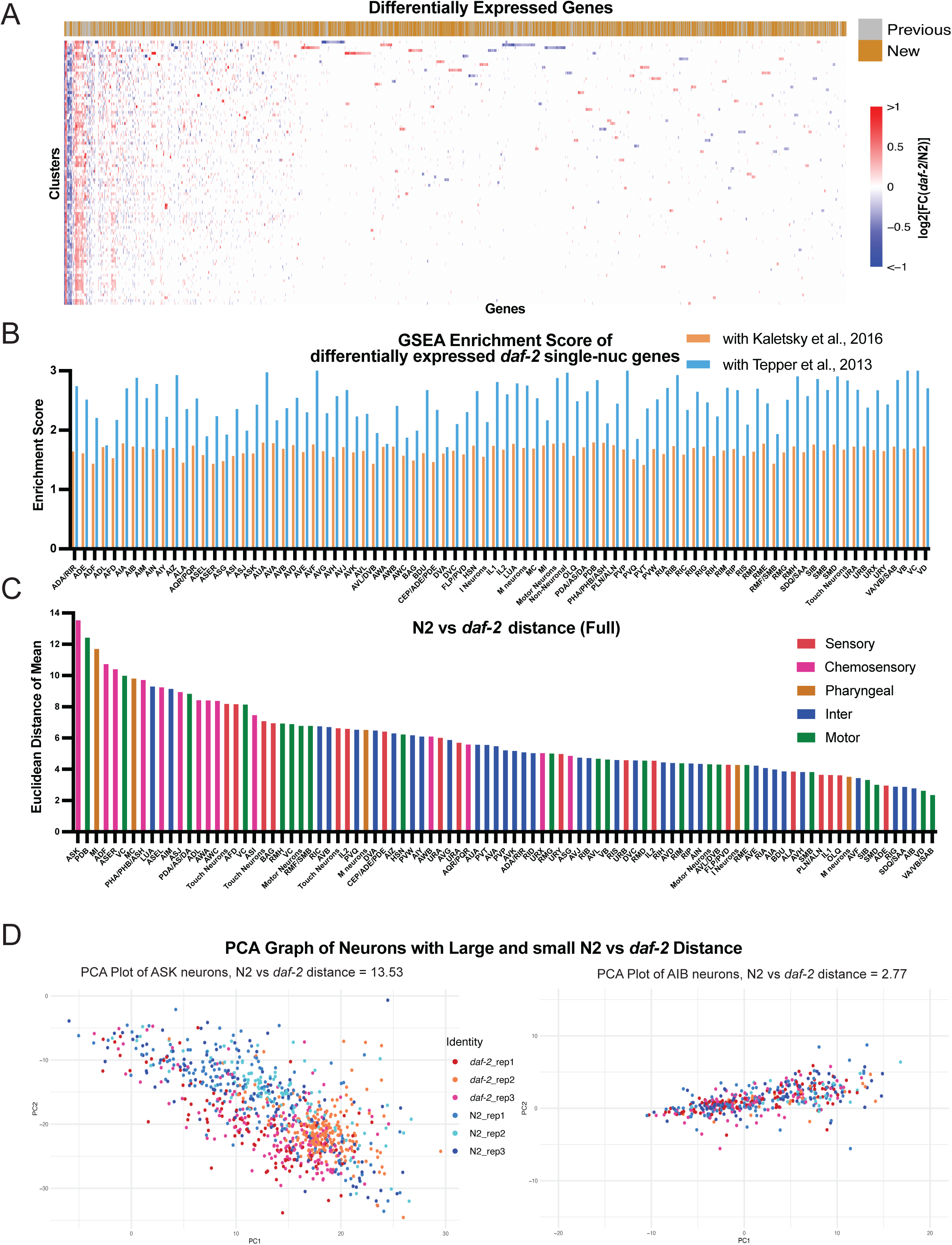
Analysis of *daf-2* vs N2 differentially expressed genes related to Figure 3. (**A**) Heatmap of differentially expressed genes across all clusters. Each row is a cluster and each column is a differentially expressed gene. Neuronal clusters and genes are hierarchically clustered. The annotation row denotes whether the gene was previously identified as a neuronal IIS target in previous studies^2,3,16^. (**B**) GSEA Enrichment score comparing the *daf-2* vs N2 differentially expressed genes in each neuron to previous bulk-sequencing results. We used GSEA to compare the gene sets generated from the *daf-2*-upregulated genes in each neuron to the ranked lists generated from *daf-2* differentially expressed genes in Kaletsky et al., 2016and Tepper et al., 2013 ^2,3^. The enrichment scores are positive for all neurons, indicating the *daf-2* vs N2 differentially expressed genes are in the same trend as previous results (**C**) Full image of N2 vs *daf-2* Euclidean distance. Related to Figure 2B. (**D**) PCA graph of individual cells across biological replicates. PCA plot of ASK neuron (N2 vs *daf-2* Euclidean distance = 13.53) and AIB (N2 vs *daf-2* Euclidean distance = 2.77) are shown as representative images. Each biological replicate is shown in different colors as indicated in the legend.

**Extended Data Figure 6.**
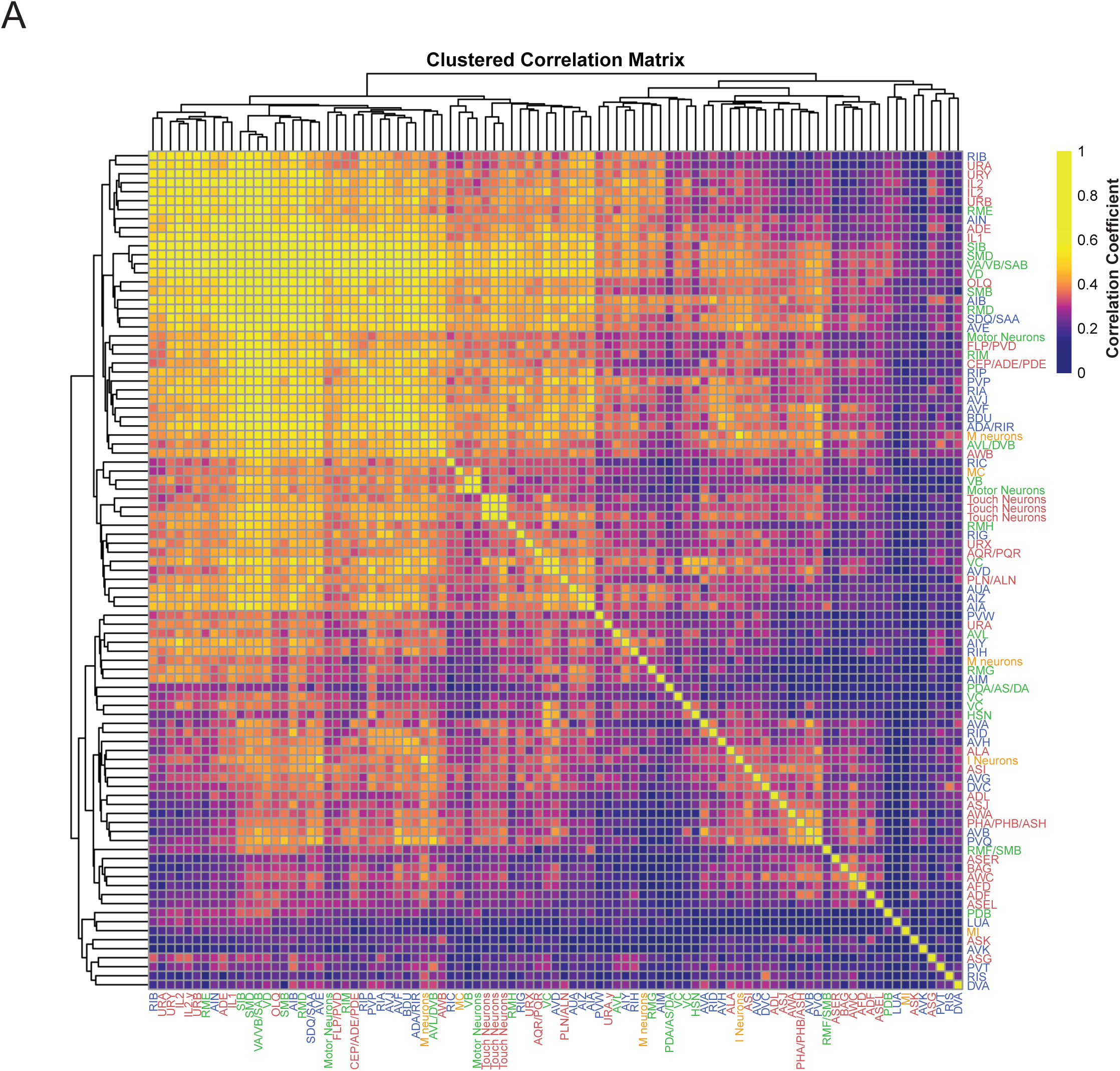
Correlation Matrix of daf-2 vs N2 differential expression. Pearson’s correlation between *daf-2* vs N2 differential expression log2(fold-change) vector was calculated between each 2 cluster pairs. The heatmap was hierarchically clustered before plotting.

**Extended Data Figure 7.**
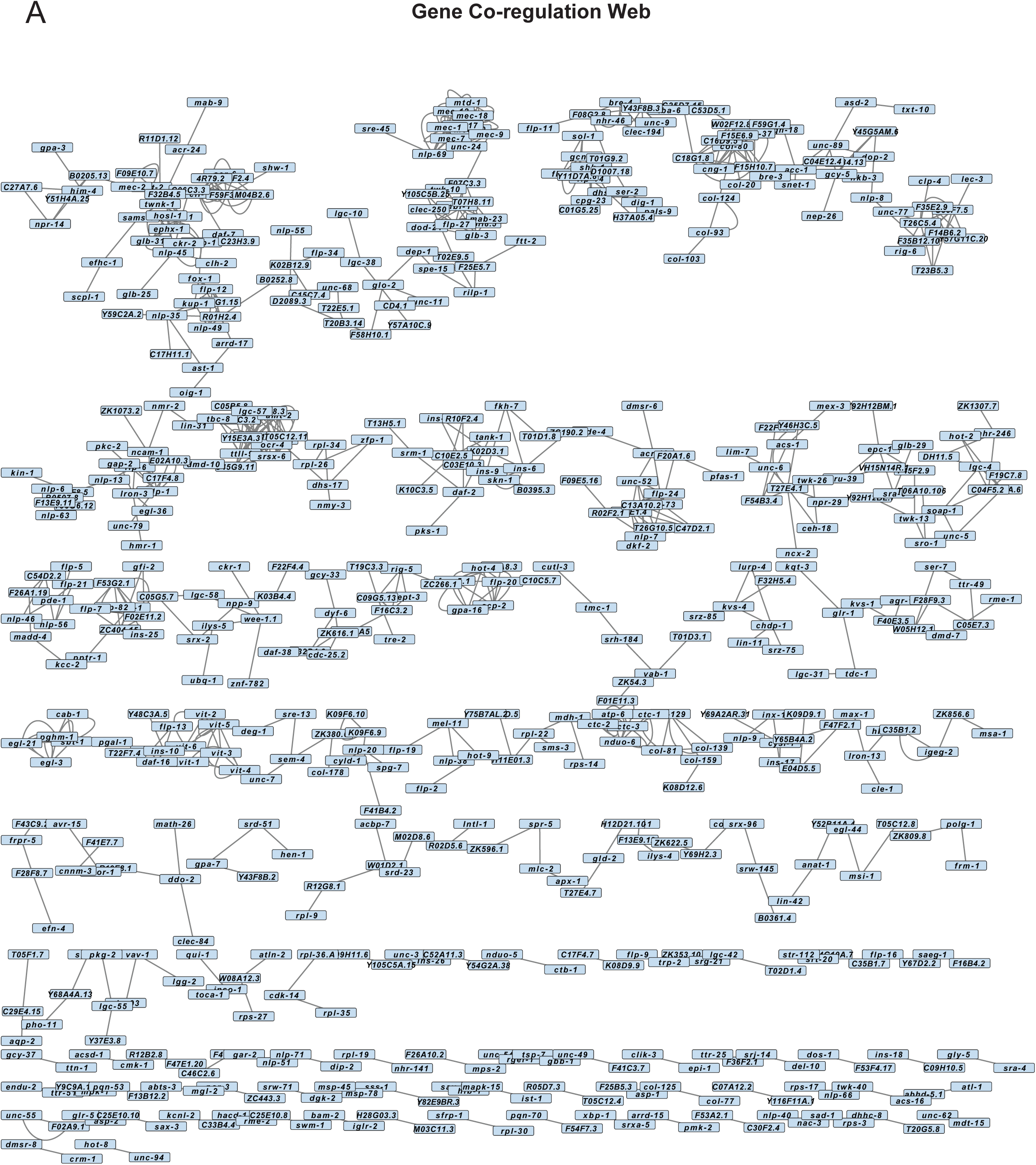
Analysis of *daf-2* vs N2 gene regulation in each cluster related to Figure 3. Full image of gene regulation web across clusters. Genes with a regulation weight greater than 0.04 calculated from GENIE3 were plotted here. The length of edges negatively correlates with the strength of co-regulation.

**Extended Data Figure 8.**
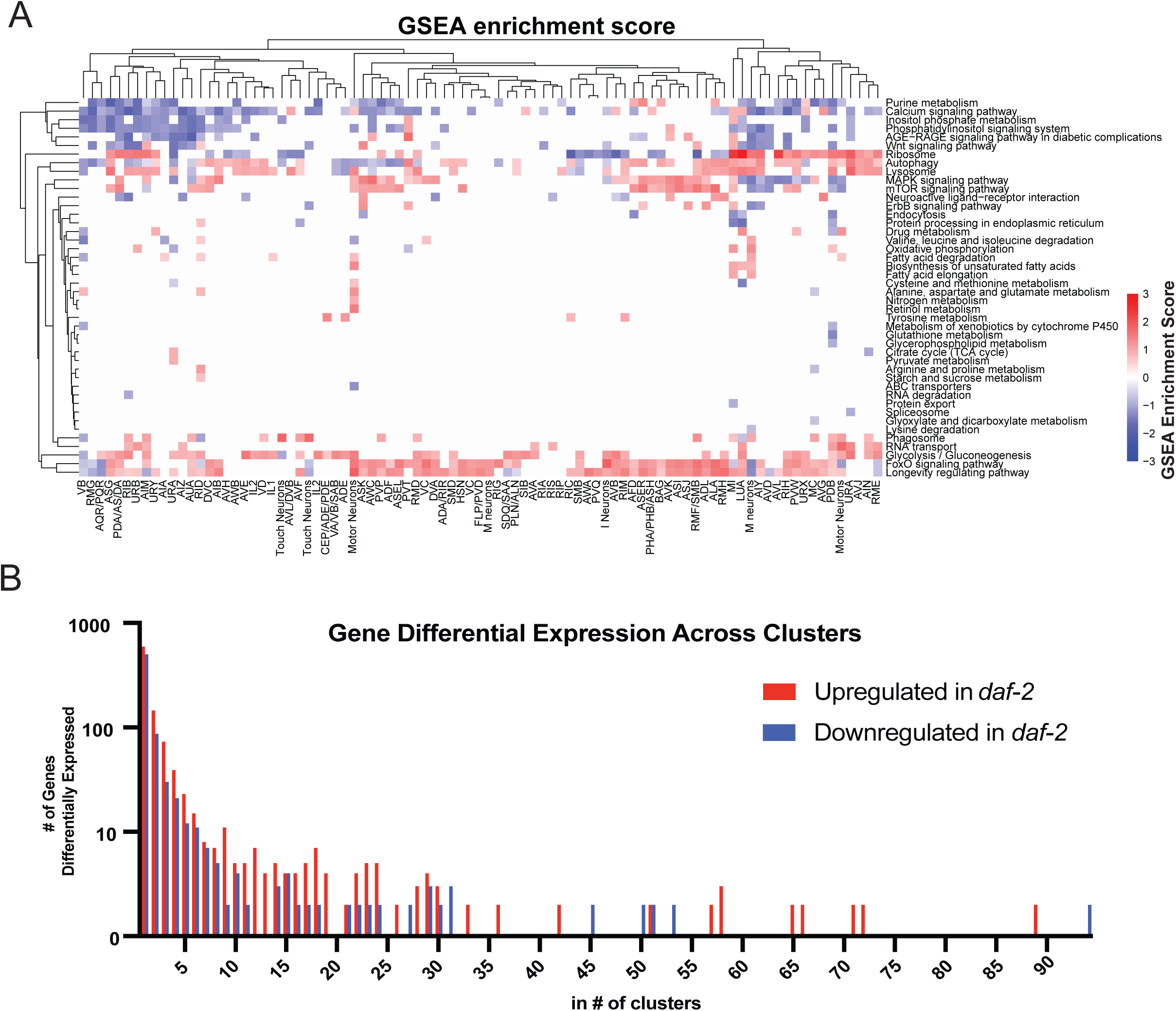
Analysis of *daf-2* vs. N2 differential expressed pathways and gene numbers, related to Figure 3. (**A**) Full image of daf-2 vs N2 KEGG pathway enrichment. GSEA analysis was performed on each cluster and normalized enrichment score was plotted as a heatmap. KEGG pathways enriched in the *daf-2* upregulated genes (red) and *daf-2* downregulated genes (blue). Only KEGG pathways with at least 1 cluster with a significant enrichment score was plotted. (**B**) Histogram of number of clusters each gene is differentially expressed in. Most genes are differentially expressed in less than 5 clusters. Very few (less than 2%) genes are differentially expressed across >30 clusters.

**Extended Data Figure 9.**
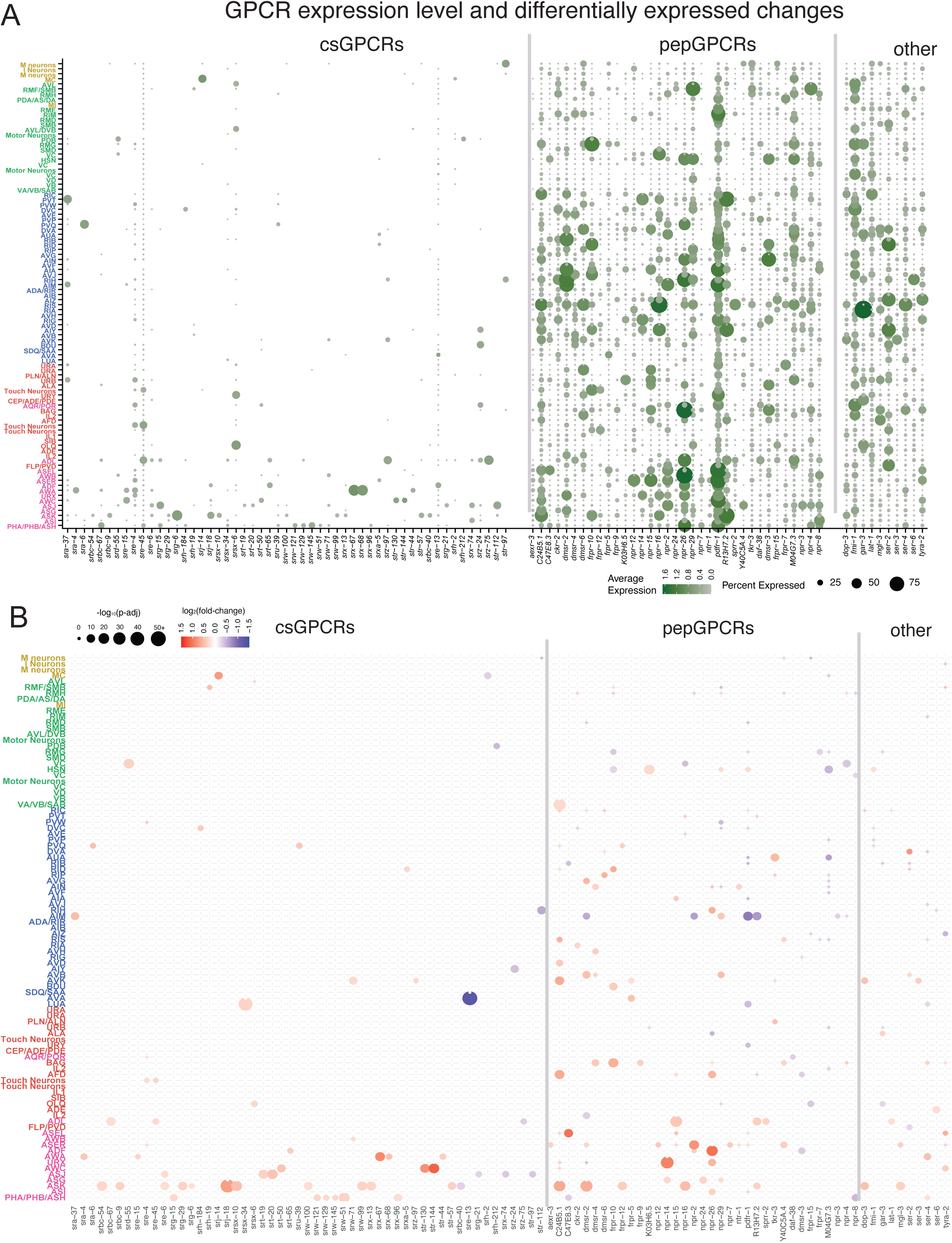
GPCR expression in combined wild-type and *daf-2* data and differential expression in wild-type vs *daf-2* neurons and their differentially expressed changes in *daf-2* neurons. (**A**) GPCR expression in different neurons of the wild type and *daf-2* combined data. The combined wild-type and *daf-2* average expression and percent expressed level of GPCRs. Only the GPCRs that are differentially expressed in *daf-2* in at least one neuron are shown. (**B**) Differentially expressed fold-change and adjusted p-values of GPCRs in daf-2 neurons. Only the GPCRs that are differentially expressed in *daf-2* in at least one neuron are shown. Related to Figure 3L.

**Extended Data Figure 10.**
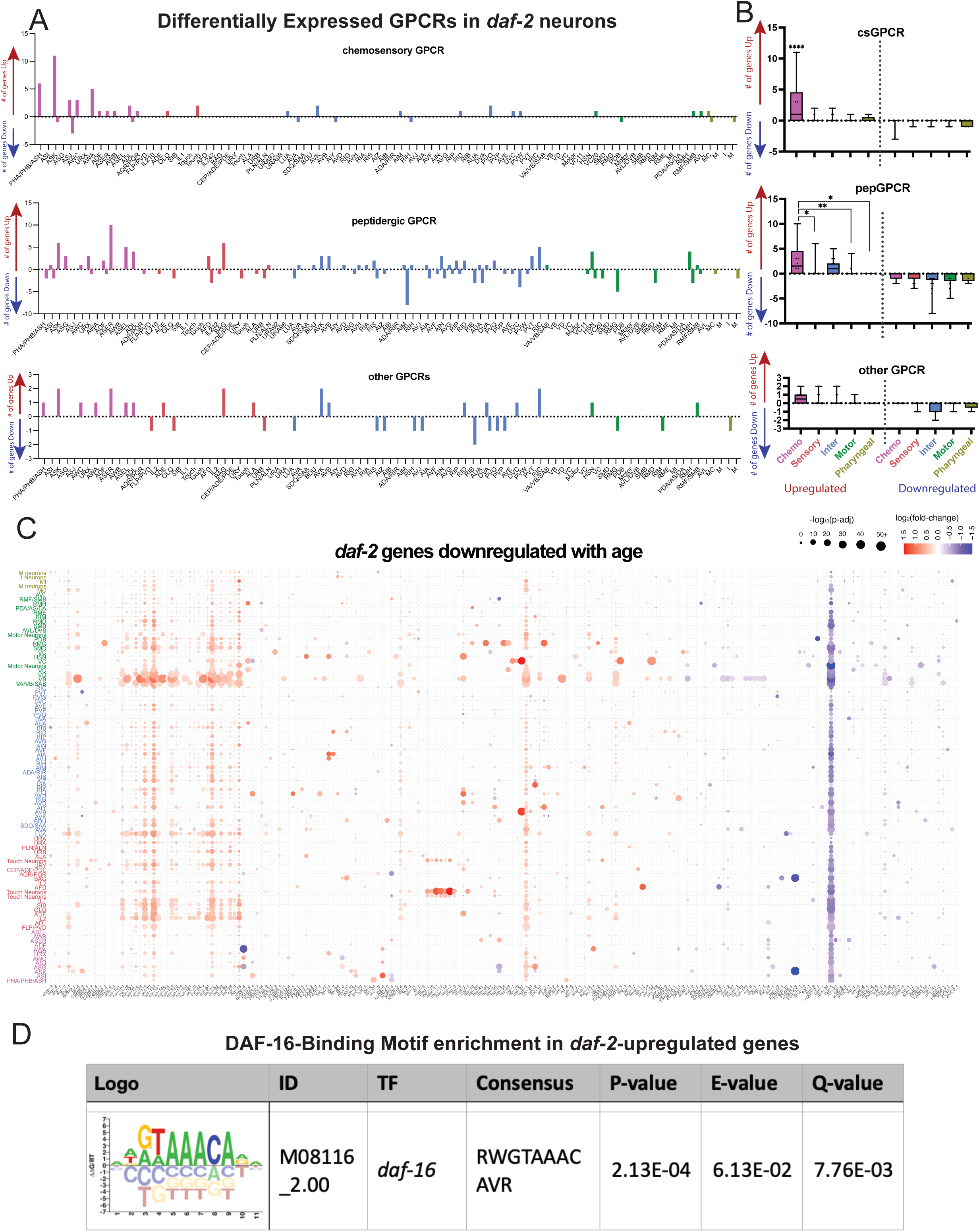
*daf-2*-Differentially expressed GPCR statistics and plot of genes that are downregulated with age. (**A**) Number of GPCRs upregulated and downregulated in *daf-2* in each neuron. csGPCRs, pepGPCRs, and other GPCRs are plotted individually. Chemosensory neurons, other sensory neurons, interneurons, motor neurons, and pharyngeal neurons are labeled in pink, red, blue, green, and yellow, respectively. (**B**) An average number of *daf-2*-upregulated and *daf-2*-downregulated GPCRs per neuron in each neuronal class. Chemosensory neurons, on average, have a greater number of chemosensory and peptidergic GPCR genes upregulated in *daf-2*. Each dot indicate a specific neuronal cluster. Box plots: center line, median; box range, 25-75^th^ percentiles; whiskers denote minimum-maximum values. One-way ANOVA. ****: p<0.0001, **: p<0.01, *: p<0.05. (**C**) Full image of Figure 4A. Single-nuc *daf-2* differential expression of genes downregulated in aged WT animals. Genes shown are those that are downregulated with age (more highly expressed in young versus old wild-type neurons) and are differentially expressed in *daf-2*.

**Extended Data Figure 11.**
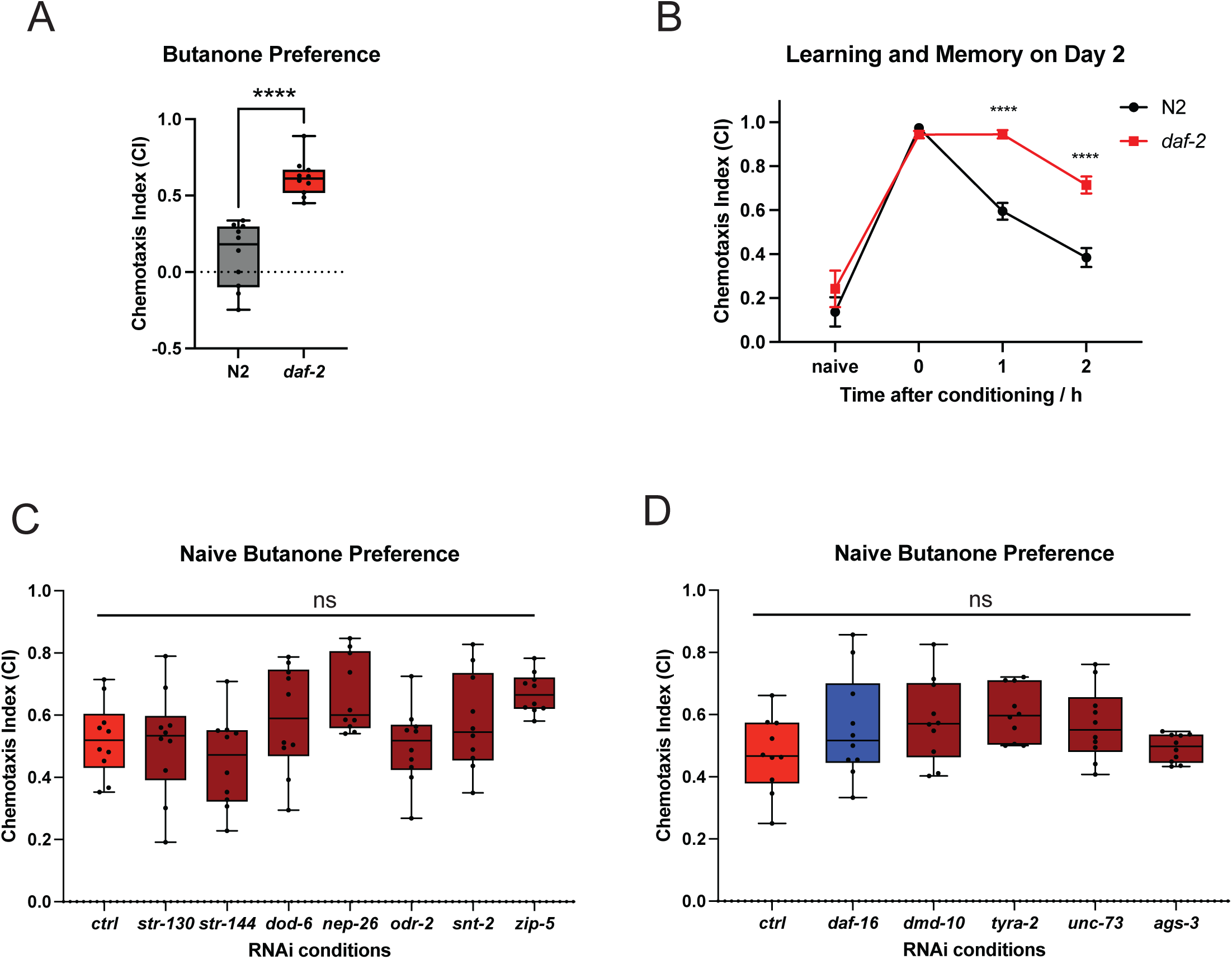
AWC-specific *daf-2* differentially expressed genes regulate cognitive behaviors in daf-2. (**A**) Naïve butanone preference is higher in *daf-2* worms. Day 1 N2 and *daf-2* worms without training were tested for their chemotaxis index towards butanone. Representative figure from 3 biological replicates. Box plots: center line, median; box range, 25-75th percentiles; whiskers denote minimum-maximum values. ***p<0.001, Student’s t-test. (**B**) Memory ability is different between N2 and *daf-2* mutants on Day 2 of adulthood. ****p<0.0001, Two-way ANOVA with Sidak’s post-hoc analysis. Error bar: SEM. (**C-D**) Naïve butanone preference in worms treated with adult-only control or RNAi targeting indicated genes. Representative figure from 2 biological replicates. Naïve butanone preference is unaffected after adult-only RNAi in Day 5 *daf-2* mutants. Box plots: center line, median; box range, 25-75th percentiles; whiskers denote minimum-maximum values. ns = not significant. One-way ANOVA. Related to Figure 5.

## Methods

### Animal growth and maintenance

All strains were maintained at 20C for the duration of our experiments. Animals were maintained on plates made with high growth medium (HGM: 3 g/L NaCl, 20 g/L Bacto-peptone, 30 g/L Bacto-agar in distilled water, with 4 mL/L cholesterol (5 mg/mL in ethanol), 1 mL/L 1M CaCl2, 1 mL/L 1M MgSO4, and 25 mL/L 1M potassium phosphate buffer (pH 6.0) added to molten agar after autoclaving). All assays were performed on plates made with standard nematode growth medium (NGM: 3 g/L NaCl, 2.5 g/L Bacto-peptone, 17 g/L Bactoagar in distilled water, with 1 mL/L cholesterol (5 mg/mL in ethanol), 1 mL/L 1M CaCl2, 1 mL/L 1M MgSO4, and 25 mL/L 1M potassium phosphate buffer (pH 6.0) added to molten agar after autoclaving^68^. All experiments that did not involve RNAi treatment were seeded with OP50 E. Coli (From the CGC) for *ad libitium* feeding. Hypochlorite synchronization was used to developmentally synchronize experimental worms, where gravid hermaphrodites were exposed to an alkaline-bleach solution (e.g. 6 mL sodium hypochlorite, 2.5 mL KOH, 41.5 mL distilled water) to collect eggs, followed by repeated washes with M9 buffer (6 g/L Na2HPO4, 3 g/L KH2PO4, 5 g/L NaCl and 1 mL/L 1M MgSO4 in distilled water)^68^. Strains used in this study: CQ757: *wqIs7 [Prgef-1::his-58::GFP]*, CQ758: *wqIs7 [Prgef-1::his-58::GFP];daf-2(e1370) III*, LC108: *vIs69 [pCFJ90(Pmyo-2::mCherry + Punc-119::sid-1)] V*, CQ745: *daf-2(e1370) III;vIs69 [pCFJ90(Pmyo-2::mCherry + Punc-119::sid-1)] V*

### Neuronal nuclei isolation

*C. elegans* neuronal nuclei were isolated using the following methods, modified from the mammalian single nucleus RNA-seq protocol from 10X Genomics. First, ∼400 μL of Day 1 worms were washed from HG plates. Worms were washed 3x in M9 buffer and transferred to a dounce homogenizer (Kimble Glass Tissue Homogenizer, 88 mm overall length, Dounce 1 mL working capacity; Cat # 885300-0001) filled with 300 μL of ice cold NP40 lysis buffer (10 mM Tris-HCl (Sigma-Aldrich, Cat. # T2194; pH 7.4), 10 mM NaCl (Sigma-Aldrich, Cat. # 59222C), 3 mM MgCl_2_ (Sigma-Aldrich, Cat. # M1028), 0.05% Nonidet P40 Substitute (Sigma-Aldrich, Cat. # 74385), 1 mM DTT (Sigma-Aldrich, Cat. # 646563), and 1 U/μL RNase inhibitor (Sigma Protector RNase inhibitor; Sigma-Aldrich, Cat. # 3335402001), Nuclease-free water). Each sample was dounce homogenized 25-35x, using only the tight pestleuntil complete worm lysis was achieved. Samples were moved to low bind microcentrifuge tubes, incubated for 5 min on ice, and pipetted 10 times at two intervals during the incubation (at 2 min and 4 min) to disrupt tissues and cells. Nuclei suspensions were passed through a 40 μm filter into a conical tube, and then transferred to a low bind microcentrifuge tube. Samples were centrifuged at 1000 x g for 5 min at 4°C. The supernatant was removed and 1 mL of wash buffer (PBS + 0.5% BSA (Miltenyi Biotec, Cat. # 130-091-376) + 1 U/uL of RNase inhibitor) was added with no mixing and incubated for 5 min on ice. After 5 min, the pellet was resuspended. Samples were centrifuged again at 1000 x g for 5 min at 4°C. The supernatant was removed, and the pellet was then resuspended in 500 μL of wash buffer. Hoechst stain (1:10,000 dilution; Molecular Probes Hoechst 33342, Thermo Fisher, Cat. # H3570) was added to each sample, and samples were passed through 5 μm syringe filters, directly into FACS tubes. Samples were incubated for at least 5 min on ice prior to FACS. Hoechst and GFP+ nuclei were sorted using a 70 μm nozzle and a flow rate of 3 on a BD Biosciences FACSAria Fusion sorter into a 1.5 mL low bind microcentrifuge tube containing collection buffer (500 μL of 0.5% BSA + 1.5. U/μL RNase Inhibitor). The instrument was washed with bleach between samples.

### snRNA-seq library preparation and sequencing

After FACS, samples were centrifuged at 1000 x g for 5 min at 4°C. The supernatant was removed and nuclei were resuspended in 20 μL of collection buffer, and nuclei integrity was monitored by microscopy, with ∼90% of nuclei intact (no membrane blebbing) for each sample. The total number of nuclei for each sample was estimated and provided to the Princeton Genomics Core. Single nuclei suspension samples were loaded to the 10X Genomics Chromium X system using the Single Cell 3’ v3.1 Reagent Kits (10X Genomics Inc., CA) to generate and amplify cDNA. Up to 30,000 nuclei were loaded per 10X sample. The amplified cDNA samples were purified with Ampure XP magnetic beads (Beckman Coulter, CA), quantified by Qubit fluorometer (Invitrogen, CA), and examined on Bioanalyzer with High Sensitivity DNA chips (Agilent, CA) for size distribution. Illumina sequencing libraries were generated from the amplified cDNA samples using the Illumina Tagment DNA Enzyme and Buffer kit (Illumina, CA). These libraries were examined by Qubit and Bioanalyzer, then pooled at equal molar amount and sequenced on Illumina NovaSeq 6000 S Prime flowcells as 28+94 nt pair-end reads following the standard protocol. Raw sequencing reads were filtered by Illumina NovaSeq Control Software and only the Pass-Filter (PF) reads were used for further analysis.

### Alignment and Quality control of data

After Illumina sequencing, alignment of reads was performed using CellRanger version 7.1.0. Quality control of the data began with removal of ambient RNA contamination using SoupX on the Cell Ranger output files. SoupX calculated contamination fractions for all samples were between 0.03 and 0.12. Next, metrics of genes/cell, average UMIs/cell, and number of cells per cluster were determined. Violin Plots of genes per cell were generated to determine cutoffs: the lower bound was 200-300 features/cell for each sample (to remove damaged nuclei and empty droplets) and the higher bound was between 1000 and 2000 features/nucleus (to remove doublets) depending on the sample. Data outside of these cutoffs were excluded from further analysis.

### Normalization, Integration, and Clustering

In Seurat, we followed the single cell pipeline, where single cell transform (SCT) was used for normalization. Prior to normalization, all data were pooled.. SCT normalization is different from the normalization used for bulk sequencing datasets: it fits genes to a negative binomial distribution to normalize them across data instead of dividing everything by a ‘size factor’ as in bulk seq. Elbow plots were generated to determine the number of principal components (PCs) to include in the dimensional reduction, and the Louvain algorithm was used for unsupervised network clustering. We tested different numbers of PCs: 80, 100, 130, 150, 200, and different resolutions: 0.5, 0.7, 1.0, and 1.2. 150 PCs at a clustering resolution of 1 was used, which resulted in 100 clusters in our dataset without overclustering of neuron subtypes.

### Cluster Labeling/ Cell Type Analysis

We performed cluster annotation using a combination of systematic and manual approaches. Initially, we utilized the ’FindAllMarkers’ function in the Seurat package to identify cluster-specific markers. These markers corresponded to genes showing significant differential expression (log2FC > 0.25) within each cluster compared to the average expression across all clusters. These markers also needed to be expressed in greater than 25 percent of cells for them to be used.

To further refine the annotation process, we compared the identified markers for each cluster with a curated list of known markers specific to each neuron type classification (Table S3). This comparison was conducted using a hypergeometric test and assessing the Bonferroni-corrected p-value for the overlapping gene regions. Clusters demonstrating a significant overlap (p < 0.01) with known anatomy markers were considered for the corresponding neuron type.

Additionally, we employed the AUCell algorithm as an independent mechanism for cluster annotation. Initially, we constructed a selected gene set collection utilizing known markers associated with each neuron type. Subsequently, we ranked the expression levels of each gene within every cell in our sequencing dataset. Using these rankings, we calculated the area-under-the-curve (AUC) value for each gene set in every cell.

To assign neuron types to cells, we generated a histogram of AUCell values and applied a threshold. In some cases, we manually adjusted the threshold to ensure approximately 5% of the cells were assigned to each neuron term. Subsequently, we assembled the cells into clusters and determined the percentage of cells within each cluster assigned to a specific neuron type. The neuron types with the highest five percentages were considered further for that cluster.

To make the final neuron type assignment for a cluster, we integrated the results from the hypergeometric test and the AUCell algorithm. If the outcomes from both methods aligned, we regarded that neuron type as the final annotation. In cases of disagreement, we manually examined gold-standard markers associated with that neuron type and made a manual decision regarding the appropriate annotation. The results of both mechanisms and the final annotation are summarized in supplemental table 3.

### Threshold Setting for Expressed Genes

When comparing our dataset to L4 data, we not only subsetted our data to only compare our wild-type data with the L4 data, we also applied expression level thresholds to the genes. We applied 4 thresholds (as done by CeNGEN^20^) that require a varying percent of expression of a gene within a cluster (0.5, 1, 1.5, or 3) and a minimum average normalized expression value (0.001). When comparing our data to CeNGEN, we always compare our 2^nd^ threshold against their 2^nd^ threshold. These thresholds show enrichment of genes in specific places and help to cut down any noise in the data.

### Hierarchical Clustering

To generate a gene-expression-based hierarchical clustering dendrogram, we first aggregated the normalized gene expression level of each gene in a cluster to obtain the average gene expression vector for each cluster, then calculated the Euclidean distance matrix between clusters, and then hierarchically clustered using the “hclust” function with the “complete” linkage method.

### Differentially-expressed gene identification

We used the FindMarkers function in the Seurat package for differential expression identification. We performed the Wilcoxon Rank Sum Test method on each subsetted cluster separately, comparing N2 cells and *daf-2* cells within the same cluster. Genes with a minimum percentage of expression in 10% of the cells were analyzed, and the significantly differentially expressed genes were identified if their log2(fold-change) > 0.25 or < -0.25, and adjusted p-value < 0.05. The genes significantly differentially expressed in at least one cluster are considered differentially expressed.

For comparison with previous datasets, genes that are differentially expressed with a p-value < 0.05 were considered differentially expressed in Kaletsky et al., 2016 dataset^2^. Genes with a ranking < 1500 or > -1500 and FDR < 0.05 were considered differentially expressed in the Tepper et al., 2013 dataset^3^. Genes that have a *daf-2* vs N2 log2(fold-change) > 0.5 and p-adjusted < 0.05 were considered differentially expressed in the Weng et al., 2023 dataset^16^. Genes present in those 3 datasets were considered previously identified and the differentially expressed genes not present in these 2 datasets were considered newly identified.

When calculating the correlation of differential expression of clusters, each cluster was represented as a vector where each dimension is the differential expression log2(fold-change) of a gene, and Pearson’s correlation of each vector was calculated to generate a correlation graph for Extended Data Figure 6A.

Gene differential expression heatmaps and dotplots were generated using the ggplot2 package in R and log2(fold-change) and p-adjusted values were extracted and plotted^69^.

### Gene co-regulation network analysis

We performed gene co-regulation network analysis using the *daf-2* vs N2 log2(fold-change) matrix. Using the GENIE3 package, we calculated the regulatory strength between all differentially expressed genes using tree-based ensemble methods ^39^. Then we selected the gene interaction of gene pairs with an interaction weight > 0.04, resulting in a total of 793 interaction pairs, and visualized their interactions using the Cytoscape software 3.10.0 ^70^. We used the perfuse force OpenCL layout where the length of an edge reflects the weight of the interaction. To plot a subset of interaction, we selected the nodes and edges to plot, then calculated the perfuse force OpenCL layout independently.

### Euclidean distance measure

We first subsetted the data into each cluster and further subsetted out N2 and *daf-2* cells using the Seurat package. For N2 and *daf-2* in each cluster, we generated a representative PC vector by averaging the column means of PCA embeddings from all cells in that cluster and genotype.

Then, we calculated the Euclidean distance between the PC vector of N2 and *daf-2* in each cluster. We ranked the distance from the largest to the smallest.

### GSEA

For calculating GSEA for KEGG pathways, we downloaded the KEGG pathways from Worm Enrichr^71^. Next, we generated a ranked list of the differentially expressed genes in each cluster according to their log2(fold-change). We applied GSEA on the ranked list for each cluster using the fgsea package in R^72^. The minimum size of the gene set to test is 3 and the maximum size is 100. Then, we plotted the normalized enrichment score for each pathway in the heatmap.

For calculating the GSEA enrichment score for the genes that downregulate with age, we input the ranked list of the differentially expressed genes in each cluster according to their log2(fold-change). Then, we generated a gene set of all genes that downregulate with age in wild-type neurons (log2FC > 2.0, p-adj < 0.001) and calculated the gene set enrichment score of each neuron for this gene set.

For calculating GSEA enrichment score for the bulk sequencing datasets, we generated the ranked list from the Kaletsky et al., dataset based on fold-change, and another ranked list from Tepper et al., dataset based on ranking^2,3^. Then we generated a gene set for each cluster using only the significantly upregulated genes. We performed GSEA of these 2 ranked lists on each gene set where the minimum size of the gene set to test is 10 and the maximum size is 1000, and calculated the normalized enrichment score.

### Gene Ontology analysis

We identified AWC-enriched genes using the FindMarkers function in the Seurat package to identify genes significantly differentially expressed in the AWC compared with the background (log2(fold-change) > 0.1, p-adjusted < 0.05). We identified the AWC *daf-2* upregulated and downregulated genes using the significantly up- and down-regulated genes from the gene differential expression analysis. We measured GO terms using the WormCat 2.0^67^ website and chose the significantly enriched pathways (Bonferroni adjusted p-value < 0.05).

### Motif Enrichment analysis

To identify the enriched transcription factor binding motifs of the *daf-2* upregulated genes, the - 1000 to -1 upstream region of the *daf-2* upregulated genes were downloaded using the RSAT. The -1000 to -1 upstream region of all *C. elegans* protein-coding genes were retrieved as the background control sequences. We then input the sequence into the Simple Enrichment Analysis (SEA) function in the Meme-suite website (https://meme-suite.org/meme/tools/sea). We used Cis-BP 2.00 C*. elegans* motif database as the input motif database. To identify enriched 7-mer oligos, we input the upstream region into the oligo-diff function in the RSAT website. The output enriched motifs and overrepresented oligomers are presented in Supplemental Table 10.

### Behavioral assays

All Short-Term Associative Memory assays (STAM) were performed as previously described^1,73^. Briefly, synchronized adult worms were washed and starved for an hour before training on plates with food and butanone (present on the plate lid) for one hour to form an associative memory.

Then worms were tested on chemotaxis plates with butanone and ethanol on opposite sites immediately after training for their learning ability or transferred to holding plates with food for 1 hour or 2 hours to subsequently test their memory ability. Naïve chemotaxis was measured for each treatment group in each experiment by testing the chemotaxis index of untrained worms.

The chemotaxis index was measured by the number of worms at the butanone location minus the number of worms at the ethanol location, divided by the total number of worms outside of the origin spot. The learning index is measured by subtracting the chemotaxis index of trained worms from the chemotaxis index of naïve worms. For chemotaxis assays, chemotaxis towards 1% butanone (Acros Organics; Cat. #332828-25ML) or 1% benzaldehyde (Millipore Sigma Cat. # B1334-100G) in ethanol was accessed using standard chemotaxis conditions (Bargmann et al., 1993). All chemotaxis assays used Day 2 adult, LC108 worms that completed two days of adult-only RNAi treatment.

For behavioral experiments with RNA interference, we used LC108 (*Punc-119::sid-1*) or CQ740 (*daf-2(e1370) III*; *vIs69 [pCFJ90(Pmyo-2::mCherry + Punc-119::sid-1)] V*) worms to increase the penetrance of RNAi to the nervous system. We performed adult-only RNAi by transferring synchronized L4 worms onto fresh RNAi plates supplemented with IPTG, carbenicillin, FUdR and additionally freshly spotted IPTG prior to worm transfer.Worms were moved to fresh RNAi plates every 2 days before performing behavior experiments. We transfer the worms to RNAi plates without FUdR 1 day before performing behavior. For *daf-2* STAM experiments, we performed the experiment on Day 5 to test effects on aging worms.

### PA14 choice assay

Choice assays were performed as described in Moore et al., 2021^74^. Overnight liquid-grown cultures of PA14 were diluted in LB to an Optical Density (OD_600_) = 1, and 25 μL of each bacterial suspension (PA14 or OP50 control) was spotted onto one side of a 60 mm NGM plate and incubated for 2 days at 25°C. Plates were moved to room temperature for 1 h before use.

Immediately before use, 1 μL of 1M sodium azide was spotted onto each bacteria spot as a worm paralytic to capture the first choice made by each animal. Day 1 adult wild-type or *daf-2* worms raised on OP50-seeded plates were collected and washed in M9. 5 μL of washed worm pellet was spotted at the bottom of the choice assay plate and incubated at room temperature for 1 hour, when all worms were fixed at either the OP50 or PA14 spot. Worms were counted and the choice index was calculated (Choice Index = (Number of worms on OP50 – Number of worms on PA14) / (Total number of worms on OP50 + PA14)).

## Acknowledgments

We thank the Christina DeCoste and the Princeton FACS Core for assistance, Jennifer Miller, Jean Arly Vomar, Wei Wang, and the Princeton Genomics Core and for their assistance, the *C. elegans* Genetics Center for strains, and members of the Murphy lab for suggestions on the manuscript. Illustrations were created using BioRender.com.

## Funding

The Simons Foundation (811235SPI) to CTM

China Scholarship Council (CSC) to YW

NSF GRFP to JSA (DGE-2039656)

National Institutes of Health grant F32 Fellowship to MES (AG079490)

National Institutes of Health Pioneer Award to CTM (NIGMS DP1GM119167)

## Author contributions

Conceptualization: RK, JSA, YW, CTM

Methodology: RK, JSA, MES

Investigation: YW, JSA, RK, MES, RSM, SZ

Visualization: JSA, YW

Funding acquisition: CTM

Project administration: CTM

Supervision: RK, CTM

Writing – original draft: YW, JSA, CTM

Writing – review & editing: YW, JSA, CTM

## Data and materials availability

Raw data from sequencing experiments have been uploaded to NCBI BioProject: PRJNA1027859. All other data are available in the main text or supplementary material.

## Supplementary Tables

**Supplemental Table 1.** Analysis of synaptically enriched genes. Comparison of 542 synaptically enriched genes^75^ across our wild-type day 1 adult data and CeNGEN’s genes detected from their supplementary data^76^. Genes detected were compared in both datasets, divided by the total 542, and multiplied by 100 to yield percent of detection. The first page of this table contains a summary of these results, and CeNGEN’s data is available on their website.

**Supplemental Table 2. Gene Set Lists Neuron CREB GNAQ.** In this table we compared our wild-type day 1 genes detected and CeNGEN’s detection against 3 datasets. The Neuronally Detected tab denotes a dataset of neuronally enriched genes ^2^. The CREB_LTAM tab denotes genes upregulated in a memory dataset collected by our lab^75^. The final GNAQ tab contains orthologs of genes upregulated upon GNAQ gain of function^77^. The first tab of this table contains a summary with information on each dataset.

**Supplemental Table 3. Cluster Identification Results.** A curated list of markers for each neuron type is listed. Enriched genes for each identified cluster are shown (pct1 = cluster of interest, pct2 = all clusters). We determined the neuron-type anatomy of each cluster based on hypergeometric test and AUCell test results. The test results and the final annotation of each cluster are shown.

**Supplemental Table 4. WT csGPCRs.** Analysis of chemoreceptor GPCR expression across cell types in our wild-type dataset. The “Summary and Info” tab contains information on what each tab in the table means as well as our thresholding method. The number of receptors depends on the threshold of detection used, and raw data should be considered along with this sheet. Throughout the manuscript we use our threshold: >1% detection in cells in the cluster and >0.001 averaged normalized expression across the cluster.

**Supplemental Table 5. WT pepGPCRs.** Analysis of neuropeptidergic GPCRs in our wild-type dataset. The “Summary and Info” tab contains information on what each tab in the table means as well as our thresholding method. The number of receptors depends on the threshold of detection used, and raw data should be considered along with this sheet. Throughout the manuscript we use our threshold: >1% detection in cells in the cluster and >0.001 averaged normalized expression across the cluster.

**Supplemental Table 6. CeNGEN Comparisons to WT.** This table highlights differences in csGPCR expression between L4 and Day 1 animals by comparing our WT data to CeNGEN’s. The “Summary and Info” tab gives information on what the differences are and which data are being compared.

**Supplemental Table 7. flps and nlps.** Analysis of *flps* and *nlps* in our full WT and *daf-2* dataset for increased resolution. This table contains a Summary and Info tab explaining all tabs in the table. The “Raw Data” tab contains detection of a gene across cells in a cluster. The “Percent Expression” tab contains the percent of cells in a cluster where a gene is detected. Next, we looked at *flps* and *nlps* that have known interactions discovered to see if we could place any in specific neurons to provide neuronal context to where the peptide is coming from. The “Known Interactors” tab contains a selection of interactors and their percent expression information; these interactors were characterized in Beets et al. 2023^12^.

**Supplemental Table 8. Receptor Presence.** Analysis of neuropeptide receptor presence in our full WT and *daf-2* dataset for increased resolution. The “Summary and Info” Tab contains information on all tabs in the spreadsheet. The “Unfiltered Cell Expression” tab contains detection of a gene across cells in a cluster. The “Percent Receptor Expression” tab contains the percent of cells in a cluster where a gene is detected. The “Avg Expression of Receptors” tab contains RNA count information of the genes, and cutoffs were made to see which neurons receptors were enriched in. Finally the “Known Interactors” tab gives localization to some interactions characterized in Beets et al. 2023^12^.

**Supplemental Table 9. *daf-2* vs N2 differentially expressed genes** *daf-2* vs. N2 differential expression results from Wilcoxon Rank Sum test of every cluster. Significantly differentially expressed genes of each cluster are shown as a separate tab with their gene name, p-value, average log2Fold-change (*daf-2*/N2), expression percentage pc1 (*daf-2*) and pc2 (N2), and adjusted p-value.

**Supplemental Table 10. *daf-2* vs N2 Neuron-specific Euclidean Distance, etc analysis.** Results from the *daf-2* vs. N2 analyses. Raw results for the distance of mean PC vector between the 2 genotypes in each cluster are listed. Gene regulation network results are listed with gene names of the 2 interacting genes and their interaction strength calculated from GENIE3 for all genes interaction weight > 0.04. The neuronal correlation matrix was shown with Pearson’s correlation coefficients between every 2-neuron cluster. Comparison of each cluster’s gene differential expression results with Kaletsky et al., 2016^2^ and Tepper et al., 2013^3^ papers using GSEA were shown, with their enrichment score, normalized enrichment score, p-value, adjusted p-value, and log2error.

**Supplemental Table 11. *daf-2* vs N2 KEGG pathways GSEA.** Results from the *daf-2* upregulated and downregulated KEGG pathway analysis. The upregulated and downregulated pathways for each individual neuron cluster was calculated using GSEA and shown as different tabs. Results include pathway name, p-value, adjusted p-value, log2error, enrichment score, normalized enrichment score, overlap size, and leading-edge genes.

**Supplemental Table 12. *daf-2-*regulated genes Motif Analysis Results.** Analysis of the AWC neurons. Lists of wild type and *daf-2* combined AWC-enriched genes and AWC-enriched Gene Ontology Terms are shown. Lists of AWC *daf-2*-up- and down-regulated Gene Ontology Terms are shown.

**Supplemental Table 13. AWC-specific *daf-2* analysis.** Motif enrichment analysis of *daf-2*-upregulated genes. List of enriched TF-binding motifs from the upstream -1000 to -1 region of *daf-2*-upregulated genes. List of enriched 7-mers from the upstream -1000 to -1 region of *daf-2*-upregulated genes

