## Supplementary material for "Adult Single-nucleus Neuronal Transcriptomes of Insulin Signaling Mutants Reveal Regulators of Behavior and Learning": Supp Figs and Table Legends

Supplementary Tables

**Supplemental Table 1.** Analysis of synaptically enriched genes. Comparison of 542 synaptically enriched genes ^6^ across our wild-type day 1 data and CeNGEN’s 1^st^ threshold of genes detected supplementary data ^2^. Genes detected were compared in both datasets, divided by the total 542, and multiplied by 100 to yield percent of detection. The first page of this table contains a summary of these results, and CeNGEN’s data is available on their website.

**Supplemental Table 2. Gene Set Lists Neuron CREB GNAQ.** In this table we compared our wild-type day 1 genes detected and CeNGEN’s 1^st^ threshold of detection against 3 datasets. The Neuronally Detected tab denotes a dataset of neuronally enriched genesa. The CREB_LTAM tab denotes genes upregulated in a memory dataset collected by our lab^6^. The final GNAQ tab contains orthologs of genes upregulated upon GNAQ gain of function^7^. The first tab of this table contains a summary with information on each dataset.

**Supplemental Table 3.** **Cluster Identification Results.** A curated list of markers for each neuron type is listed. Enriched genes for each identified cluster are shown (pct1 = cluster of interest, pct2 = all clusters). We determined the neuron-type anatomy of each cluster based on hypergeometric test and AUCell test results. The test results and the final annotation of each cluster are shown.

**Supplemental Table 4. WT csGPCRs.** Analysis of chemoreceptor GPCR expression across cell types in our wild-type dataset. The “Summary and Info” tab contains information on what each tab in the table means as well as our thresholding method. The number of receptors depends on the threshold of detection used, and raw data should be considered along with this sheet. Throughout the manuscript we use our semi-liberal threshold (2) : >1% detection in cells in the cluster and >0.001 averaged normalized expression across the cluster.

**Supplemental Table 5. WT pepGPCRs.** Analysis of neuropeptidergic GPCRs in our wild-type dataset. The “Summary and Info” tab contains information on what each tab in the table means as well as our thresholding method. The number of receptors depends on the threshold of detection used, and raw data should be considered along with this sheet. Throughout the manuscript we use our semi-liberal threshold (2) : >1% detection in cells in the cluster and >0.001 averaged normalized expression across the cluster.

**Supplemental Table 6. CeNGEN Comparisons to WT.** This table highlights differences in csGPCR expression between L4 and Day 1 animals by comparing our 2^nd^ threshold of expression in our WT data to CeNGEN’s 2^nd^ threshold of expression. The “Summary and Info” tab gives information on what the differences are and which data are being compared.
